# Structural basis for co-translational assembly of homo-oligomeric proteins in *cis* and in *trans*

**DOI:** 10.1101/2025.11.12.687954

**Authors:** Jiří Koubek, Sebastian Filbeck, Gretchen Fixen, Sophie Kopetschke, Jaro Schmitt, Stefan Pfeffer, Günter Kramer, Bernd Bukau

## Abstract

The efficient formation of protein complexes is essential for cellular function. Across all kingdoms of life, protein complexes frequently assemble co-translationally, either when one subunit is fully synthesized before interaction with nascent partner subunits (co-post) or when multiple nascent subunits interact co-translationally (co-co), with homomeric proteins particularly enriched among co-co assembling complexes. However, it is unknown whether co-co assembly of homomers occurs on the same mRNA in *cis* or across neighbouring mRNAs in *trans*, and how ribosomes spatially organize to enable co-co assembly. Using *E. coli* homodimeric protein PheA as a proof-of-concept model, we show by ribosome profiling and cryo-EM structural analysis that it employs the co-co assembly route. Co-co assembly of PheA is facilitated by the proximity of polypeptide exit tunnels, but does not rely on fixed ribosomal orientations. Surprisingly, PheA was identified as an exceptional candidate where *trans*-assembly is potentially the primary mode of PheA co-co assembly, generating large polysomal networks for PheA synthesis. *Cis*-assembly of PheA was less frequent and occurred mainly between non-adjacent ribosomes on the mRNA due to spatial restraints imposed by the arrangement of directly neighbouring ribosomes in polysomes. These findings reveal fundamental principles of how cells structurally organise fast and efficient co-translational protein complex assembly.

## Introduction

Most proteins function as multi-subunit complexes (*1*, *2*), requiring precise and timely assembly to ensure cellular function (*3*). Recent studies have shown that complex formation can already begin while at least one subunit is still being synthesized by the ribosome (*4–11*). This mechanism is conserved from bacteria to humans. Coupling translation with complex assembly improves the formation of protein complexes in the crowded cytosolic environment of cells, by preventing aggregation (*12*, *13*) and premature degradation of unassembled subunits (*14*, *15*), facilitating the hierarchical formation of multi-subunit assemblies (*16–18*), and expanding the range of feasible protein structures through increased folding space (*19–21*).

Two co-translational assembly modes have been identified. In co-post assembly, one subunit is fully synthesized and folded before interacting with the nascent partner (*4–6*, *8–10*), enabled by diffusion-driven interactions and frequently by proximal synthesis of interacting subunits in the cytosol. In contrast, co-co assembly involves two or more interacting nascent chains (*7*, *11*) which poses a unique challenge for nascent chains: finding the right interaction partner while still being constrained in mobility by the ribosome and mRNA (*22*).

A prevailing view holds that co-co assembly of homomeric complexes primarily occurs in *cis*, between adjacent ribosomes translating the same mRNA (*11*, *23*). However, structural studies have shown that the ribosome arrangement in polysomes spatially separates polypeptide exit tunnels to prevent misfolding and aggregation (*24–28*), thereby limiting direct interactions between nascent chains during co-co assembly. Thus, while high local ribosome density on mRNAs can increase the effective concentration of interaction-competent chains, it remains unclear how efficient co-co assembly in *cis* can be achieved in this context.

Co-co assembly in *trans*, that is between nascent chains synthesized from different mRNAs, is a prerequisite for co-translational heteromer formation (*7*). This mode of co-co assembly requires precise spatial alignment of mRNA to allow encounters between nascent chains from separate polysomes during ongoing translation. However, the structural organization that facilitates co-co assembly in *trans* and the role of relative ribosome arrangements in promoting co-co assembly in this process remain unknown.

In light of these open questions, we investigated the structural basis of co-co assembly, using the *E. coli* homodimeric enzyme PheA as a model. By combining ribosome profiling, cryo-electron microscopy and tomography (cryo-EM and cryo-ET), we dissected how ribosomes organize during co-co assembly. Our results reveal that spatial proximity of polypeptide exit tunnels is a hallmark of PheA co-co assembly. Interestingly, for PheA, *cis* assembly is relatively rare, and instead, PheA assembly predominantly involves interactions between adjacent ribosomes translating different mRNAs. These findings provide a structural view of co-co assembly in cells and reveal that *trans* assembly can be an oligomerization pathway even for homomeric complexes. Collectively, our study establishes a conceptual framework for understanding how cells organize protein complex formation.

## Results

### PheA undergoes rapid co-co assembly in *trans* after full exposure of its dimerization domain

Our previous study has shown that co-co assembly in *cis* may be an important pathway to ensure allele specificity during complex assembly (*11*). However, our information about co-co assembly in *trans* is limited, in particular to whether it exists at all and how important it is in protein complex formation. Therefore, we devised an experiment to systematically screen for co-co assembly in *trans* in wild-type cells (WT) using ribosome profiling of very heavy polysomes. Our rationale was that extensive *trans*-interactions of nascent chains will interconnect multiple mRNAs, generating large ribosome assemblies that could sediment into the pellet of the sucrose gradients used for polysome profiling. We therefore expected that these interconnected mRNAs would be recovered from the gradient pellet by proteinase K digestion of the interacting nascent chains prior to polysome profiling (**Figure 1A**). To test this hypothesis experimentally, we first confirmed that the pellet fraction of sucrose gradients contained intact ribosomes (**Figure S1**). Total translatomes generated by ribosome profiling revealed that the pelleted ribosomes were actively engaged in translation (**Figure S2**). Polysome profiles showed that the sedimentation of most mRNAs was unaffected by proteinase K treatment, suggesting that their pelleting did not depend on interactions between nascent chains. However, closer analysis of the ribosome profiling data identified 8 mRNAs that behaved differently, most prominently the mRNA encoding PheA (**Figure S3**). These results suggest that *trans* assembly may be employed by only a small fraction of co-co assembling nascent chains, including PheA

**Figure 1:**
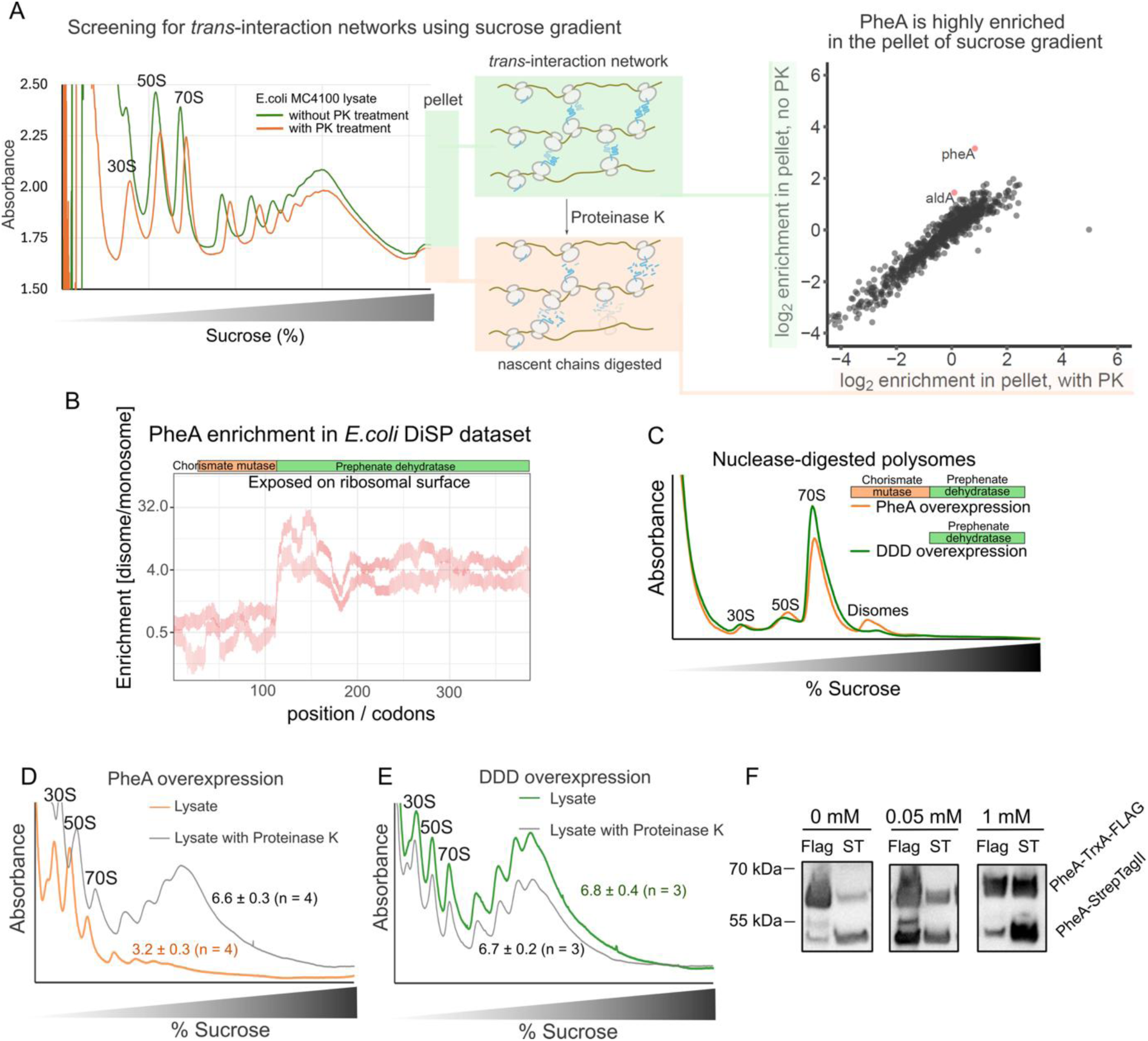
Bacterial PheA assembles co-translationally via interaction of nascent chains translated from different mRNAs. A) General experiment design with a sucrose gradient of *E. coli* MC4100 cells (left) and results of polysome profile-based screening for *trans*-interactors (right). Interaction networks are expected to shift from the pellet of sucrose gradients into the polysome fractions in a nascent chain-dependent, proteinase-K sensitive manner. Enrichment plot of several potential *trans*-assembly candidates (right). Each dot represents a translated open reading frame. The enrichment value was determined by comparing ribosome-protected footprints in the sucrose gradient pellet to total translatome, either without proteinase K treatment (y-axis) or following mild proteinase K digestion (x-axis). Red color indicates at least 2-fold depletion in the pellet after treatment with proteinase K and at least 2-fold enrichment in the pellet compared to total translatome. B) The disome enrichment profile of PheA in disome-selective profiling (DiSP). During the progression of translation, PheA footprint reads shift from monosome to disome after the exposure of the dimerization domain. The boundary of chorismite mutase domain (dimerization domain) exposed on ribosomal surface (cartoon above) is corrected for the polypeptide exit tunnel C) Overexpression of full-length wild type PheA but not of the dimerization domain deletion (DDD) PheA leads to an increased nuclease-resistant disome peak in sucrose gradients. D-E) Polysome profiles of *E. coli* overexpressing wild type PheA (D) or DDD before and after treatment with proteinase K. The ratio of area under curve of polysomes over monosome is indicated (± SEM). F) Results of pull down of double ORF experiment. PheA-StrepTagII and PheA-TrxA-FLAG tag were cloned into pET duet vector under independent promoters and cells were induced with 0 mM, 0.05 mM, or 1 mM IPTG. Proteins were pulled on from the lysate by either using Streptactin beads (ST) or anti-FLAG beads (Flag). In each condition, we observe the co-purification of the other protein variant. 0.5, 0.125, and 0.03 volumes were loaded for 0 mM, 0.05 mM, and 1 mM sample, respectively.

To confirm that these candidates are connected to co-co assembly, we analyzed disome-selective profiling (DiSP) data of WT *E. coli*. In DiSP experiments, ribosome pairs linked by nascent chains are purified from cell lysates, and nascent chains are identified through sequencing the 30 nucleotide-long ribosome footprints protected from non-collided monosomes (*7*, *11*). Among the 8 candidates, 5 showed the typical signature of co-co assembly candidates, including PheA. PheA is a 386-residue bifunctional enzyme, consisting of an 89-residue N-terminal chorismate mutase domain which contributes the main inter-subunit contacts for dimer formation (*29*). This domains chorismite mutase activity strictly requires homodimerization (*29*), and thus allowed us to differentiate between specific interactions and non-specific associations of hydrophobic nascent chains by monitoring its enzymatic activity. In addition, *pheA*-translating ribosomes rapidly shift from monosomes to disomes within a translation window of just ten codons, immediately after the full exposure of the PheA dimerization domain at the tunnel exit (**Figure 1B**). Consistent with direct nascent chain interactions, *pheA* overexpression generated a pronounced MNase-resistant disome peak that disappeared when the dimerization domain was deleted (PheA_96-386_, referred to as “dimerization domain deletion”, DDD) or mutated, or when mild proteinase K treatment was applied to digest the nascent chains (**Figure 1C**, **S4A–D**). The retained chorismate mutase activity of these disomes (**Figure S4D**) confirmed that the interactions generate functional PheA dimers during ongoing translation rather than nonspecific aggregates.

Notably, PheA overexpression led to a significant depletion of heavier polysome species (trisomes and above) in the sucrose gradient (**Figure 1D**), while intact ribosomes were enriched in the pellet (**Figure S1**). This effect was reversed by proteinase K digestion of the crude lysate and absent for dimerization-deficient mutants (**Figure 1D-E**, **Figure S4D**). These features support the model that dimerizing PheA nascent chains form extensive networks of interacting polysomes, connecting multiple ribosomes into large assemblies that sediment.

Having established that PheA undergoes co-co assembly, we next wanted to directly test whether PheA homodimers enclose subunits derived from different mRNAs. To this end, we designed a dual pull-down assay using a plasmid encoding two variants of *pheA* on separate mRNAs (**Figure S5**). One variant of PheA was C-terminally Strep-tagged (PheA-Strep) whereas the other was C-terminally tagged with thioredoxin and FLAG (PheA-FLAG). This design enabled subunit-specific purification of the two PheA constructs. Because the respective fusion proteins differ in molecular mass, their co-purification could be robustly assessed by standard SDS–PAGE. To test whether PheA undergoes *trans* dimer formation, we prepared lysates from cells grown with or without the inducer IPTG, cleared them from protein aggregates by extensive centrifugation, and performed FLAG or Strep pull-down assays. We observed that the pull-down of PheA-FLAG co-purified PheA-Strep and vice versa, providing direct evidence that PheA dimers can form from products from different mRNA transcripts (**Figure 1F**, **S6**). Notably, co-purification was not only observed under conditions of overexpression but also at basal expression levels, indicating that *trans* co-co assembly of PheA occurs under physiological expression conditions and is not merely an artifact of very high mRNA or nascent protein concentrations.

Together, these findings indicate that homodimeric PheA can assemble in *trans* by tethering multiple translating ribosomes located on different mRNAs into dynamic, higher-order polysome networks. While our data indicate that extensive cross mRNA interactions are rather rare, and in *E. coli* limited mostly to *pheA*, this finding uncovers a new layer of molecular spatial coordination in the crowded bacterial cytoplasm and provides a unique opportunity to study mechanisms of co-co assembly.

### Ribosomal polypeptide exit tunnels align in close proximity during co-co assembly of PheA

The steep disome enrichment profile in the DiSP signal (**Figure 1B**) immediately upon the synthesis of residue 115-125, corresponding to the exposure of the residue 85-95 PheA on the ribosomal surface implies that successful co-co assembly involves close spatial alignment of the polypeptide exit tunnels of two translating ribosomes. To test this directly, we visualized ribosome–ribosome arrangements during PheA co-co assembly using cryo-EM single particle analysis (cryo-EM SPA). Considering that *trans* assembly was independent of *pheA* expression levels, ribosomes were isolated from *E. coli* cells overexpressing PheA, in which 85–90% of translating ribosomes were engaged within the *pheA* ORF (**Figure S7A**). The DiSP profile confirmed that the timing and efficiency of co-co assembly in the overexpression system mirrors the endogenous context (**Figure S7B** and (*11*)), validating this approach for mechanistic analysis.

We purified monosome and disome fractions from MNase-digested polysomes and subjected them to cryo-EM SPA (**Figure 2A**, **S8A-D**, **Supplementary Table 1**). Measurement of the tunnel-to-tunnel (TtT) distance, i.e. the closest distance between polypeptide tunnel exits of adjacent ribosomes, revealed a striking difference: while monosomes showed a peak TtT distance of ∼220 Å, disomes shifted to a much shorter distance at ∼120 Å (**Figure 2B**,**C**, **S8E**, **S9A**). The small fraction of ribosomes with TtT distance of ∼ 120 Å in the monosome sample most likely originated from imperfect separation of monosomes and disomes on the sucrose gradients. As a control, randomizing ribosome orientations equalized TtT distributions across both samples (∼210–220 Å), confirming that the observed proximity in disomes reflects the preferential orientation of exit tunnels for nascent interactions during PheA dimerization (**Figure 2C**).

**Figure 2:**
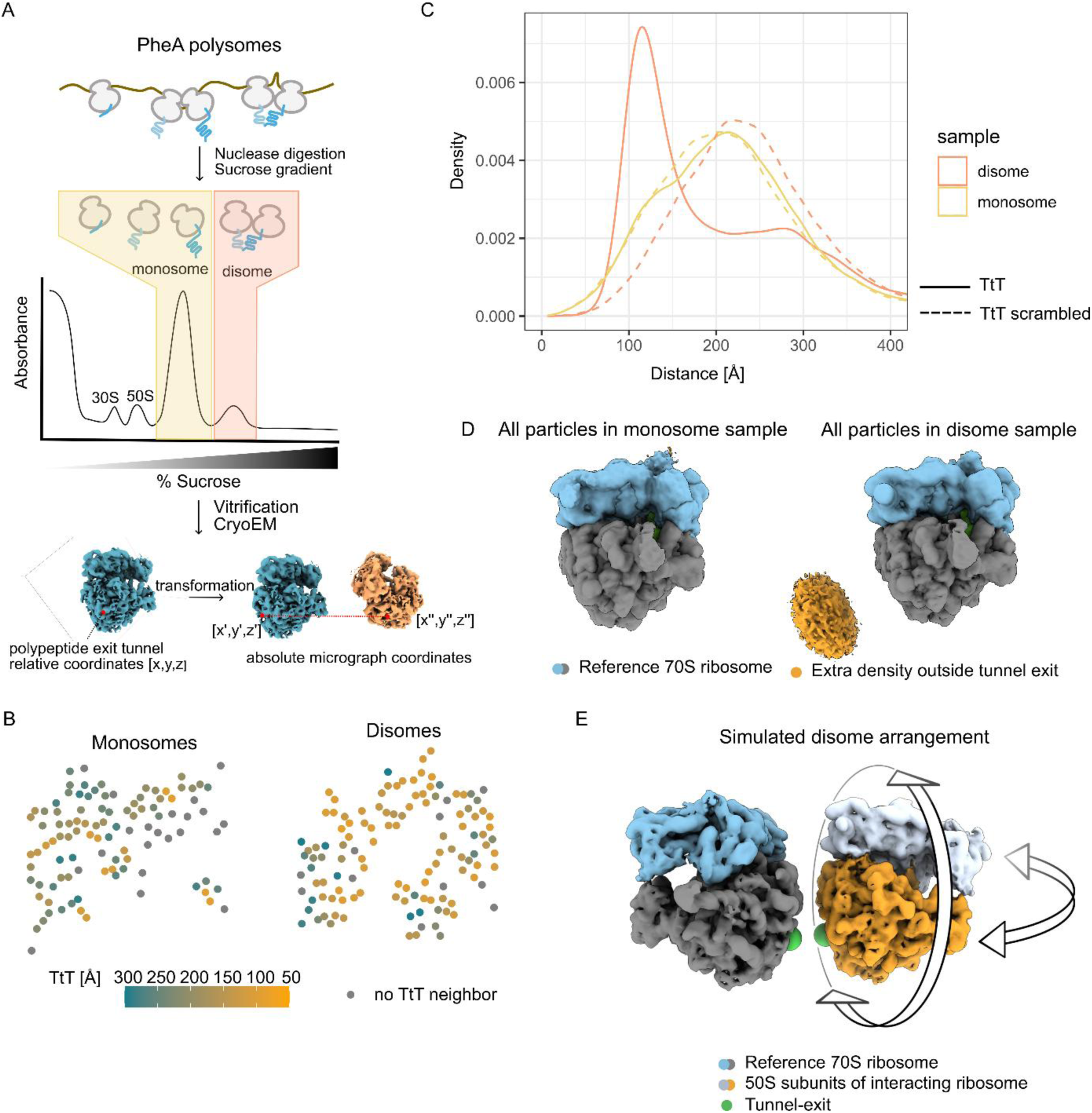
Co-co assembly coincides with polypeptide exit tunnels coming into proximity without any restriction to relative position between ribosomes. A) General processing scheme for SPA cryo-EM analysis. Lysates overexpressing PheA were resolved on sucrose gradients and fractions corresponding to monosomes and disomes were vitrified on cryo-EM grids. For each particle, relative polypeptide exit tunnel coordinates were transformed into absolute micrograph coordinates and the closest distances between polypeptide exit tunnels (tunnel-to-tunnel, TtT) were recorded. Alternatively, angles of particles were randomly scrambled and TtT distances were recorded (scrambled TtT). B) Overview of representative micrographs of monosome (left) and disome (right) samples. Each dot represents a ribosome colored according to the closest TtT neighbor distance (color code below) C) Distribution of TtT (full line) and scrambled TtT (dashed line) distances for all particles in monosome (yellow) and disome (orange) samples. D) Cryo-EM density of monosomes (left) and disomes (right) resolved to 8.6 Å resolution (50S = grey, 30S = blue). Only the disome sample contains an additional low-resolution density near the polypeptide exit tunnel (orange). E) Simulated disome arrangement. While the reference ribosomes (50S = grey, 30S = blue) are kept in the same position and orientation, interacting ribosomes (white and orange) show a high degree of orientational freedom. Polypeptide exit tunnels are indicated by a green sphere.

Consistently, cryo-EM reconstructions of MNase-resistant disomes, but not monosomes, showed a prominent additional low-resolution density positioned at the polypeptide exit tunnel (**Figure 2D**). When reconstructing only particles with TtT < 140 Å, this density became even clearer (**Figure S9B**). Focused particle extraction and subsequent refinement of this region resolved a second ribosome, also exhibiting comparable tunnel-associated density (**Figure S9C**), consistent with two ribosomes bridged by dimerizing PheA nascent chains.

To assess whether there was any preferential orientation between two interacting ribosomes, possibly arising from specific interactions between ribosomes or the dimerizing nascent chains, we applied multidimensional cluster analysis (next neighbour distribution, NND, (*25*)) and sorted interacting ribosome pairs according to their spatial and angular configurations (**Figure S9D**, **S10**). Ribosome pairs sampled all sterically allowed configurations while maintaining close TtT distances (**Figure 2E**), suggesting that ribosomes interact in a highly flexible way through the dimerizing nascent chains.

Together, these results demonstrate that during PheA co-co assembly, ribosomes align their polypeptide exit tunnels within ∼120 Å, enabling direct interaction of short, emerging nascent chains. This proximity acts as a structural signature of co-co assembly, while the flexible geometry of ribosome pairs suggests that the principal requirement is exit tunnel alignment, with no specific mRNA-independent higher-order ribosome arrangement imposed.

### Tunnel-to-Tunnel proximities are preserved in purified PheA polysomes and occur mostly in *trans*

Previous structural studies suggested that ribosome arrangement in polysomes may spatially separate polypeptide exit tunnels of adjacent ribosomes, limiting opportunities for nascent chain interactions (*24–28*). To test if and how this impacts PheA co-co assembly, we performed cryo-EM SPA on intact polysomes isolated from WT *E. coli* (MC4100) and cells overexpressing either PheA (PheA) or a dimerization domain deletion mutant (DDD) (**Figure S11**, **Supplementary Table 1**). We traced polysomes by mapping the 3D coordinates of mRNA entry and exit sites between adjacent ribosomes and assigned ribosome positions along each mRNA (*28*, *30*). Additionally, we registered each ribosome’s closest polypeptide exit tunnel neighbour (**Figure 3A**).

**Figure 3:**
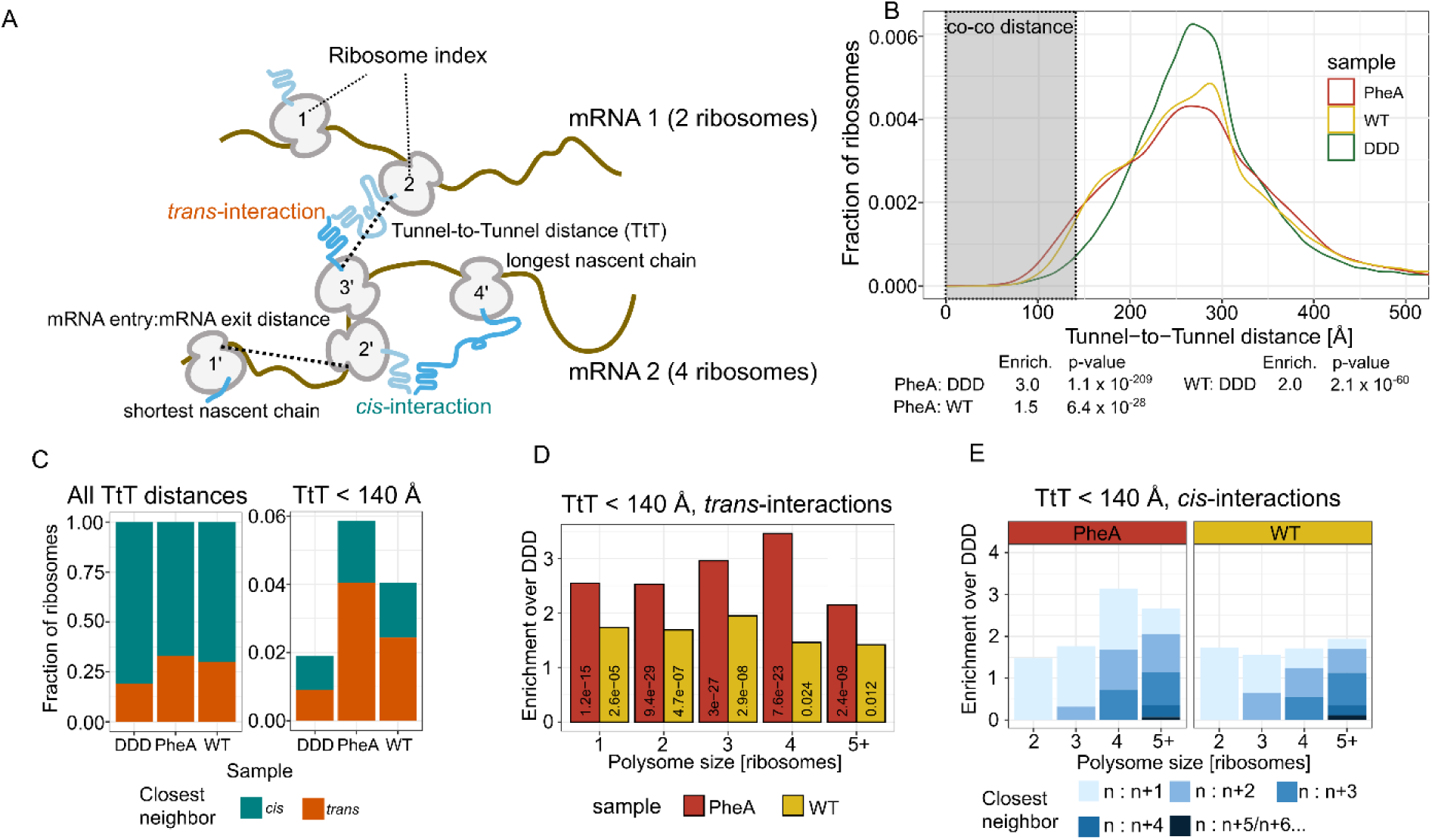
Polypeptide exit tunnels come into proximity in purified polysomes during co-co assembly. A) General overview of features measured by spatial and geometric polysome analysis. Based on TtT and mRNA entry:mRNA exit distances, each localized particle is assigned to a polysome, relative position within a polysome, and identity of its closest polypeptide exit tunnel neighbor. B) Overall TtT distributions in PheA, WT and DDD samples. Coco distance enrichment and p-values of Fisher’s exact test are indicated under the plot. C) Assignment of closest TtT neighbors on either same mRNA (*cis*) or different mRNA (*trans*) across all distances (left) or co-co distances (right). D) Enrichments of *trans*-interactions in PheA (red) and WT (yellow) samples over DDD, analyzed by the assigned polysome size. Fisher’s exact test p-values are indicated within each bar. E) Enrichments of *cis*-interactions in PheA (left panel) and WT (right panel) samples over DDD, analyzed by the assigned polysome size. The distribution of relative positions between closest neighbors (interaction partners) is indicated by the color within each bar.

Short-range TtT distances (0–140 Å) matching those of MNase-resistant disomes, interacting via their nascent chains (**Figure 2B**) were detected in all samples. However, these short-range distances were enriched 3-fold in PheA-overexpressing cells compared to DDD controls (**Figure 3B**), demonstrating that close tunnel proximity is preserved in intact polysomes and marks productive co-co assembly. WT cells also showed a ∼2-fold enrichment over DDD, suggesting that co-co assembly is widespread in the *E. coli* proteome under physiological conditions, without PheA overexpression (**Figure 3B**; **Figure S12**). Low levels of short TtT distances were also observed in DDD samples and likely reflect random polypeptide exit tunnel proximities on the EM grid. Importantly, comparable enrichment of short TtT distances in PheA overexpressing and WT samples over samples of DDD overexpressing cells was robustly observed over a range of different TtT cut-offs (**Figure S13**).

Classifying tunnel neighbours by mRNA assignment revealed that the vast majority of short TtT proximities in PheA polysomes (from PheA overexpressing samples) arose in *trans*, between ribosomes translating different mRNAs (**Figure 3C**) (so-called *trans* contacts), consistent with our polysome profiling data (**Figure 1**). We observed a similar trend in WT samples suggesting that *trans* co-co assembly is also possible for endogenous nascent chains.

To test whether ribosome density on mRNAs affects *trans* co-co assembly, we determined the frequency of *trans* contacts between ribosomes on polysomes assigned with different numbers of ribosomes. For polysomes with detectable 2 to >5 ribosomes and even monosomes, the *trans* contacts were similarly (>2-fold) enriched in PheA compared to DDD (**Figure 3D**), indicating that *trans* assembly persists across a range of translation initiation rates.

Together, these analyses show that close polypeptide exit tunnel proximity, indicative of nascent chain interaction, is maintained in intact polysomes isolated from cells and for PheA primarily occurs in *trans*, forming large, connected ribosome networks during translation.

### TtT proximity in *cis* is rare and preferentially occurs between distant ribosomes in PheA polysomes

Although *trans* contacts dominate for PheA polysomes, we also detected short-range TtT contacts within the same mRNA (so-called *cis* contacts). These *cis* contacts were more common in PheA than in DDD samples but remained far less abundant than *trans* contacts at distances < 140 Å (**Figure 3C**).

Previous work predicted that *cis* co-co assembly should require high ribosome density to increase the chance that adjacent ribosomes align productively (*11*). Consistently, *cis* contacts were enriched only in longer polysomes (≥ 4 ribosomes), whereas *trans* contacts remained enriched across all polysome sizes (**Figure 3D**,**E**).

Examining ribosome positions revealed that unexpectedly, *cis* contacts typically occurred between ribosomes separated by one to three other ribosomes (n+2 or higher) on the same mRNA (**Figure 3E**). In densely packed polysomes, polypeptide tunnel exits were observed to be spatially separated in directly neighbouring polysomal ribosomes (*24–28*). Consistently, we observed that short mRNA exit-to-entry distances were associated with long TtT distances, and vice versa, in directly neighbouring ribosomes (**Figure S14**). Thus, polysome looping may allow distant ribosomes in densely packed polysomes to orient in a configuration that enables polypeptide exit tunnel to come in close proximity.

Collectively, these findings indicate that *ci*s co-co assembly is rare for PheA, requiring long polysomes and interactions between distal ribosomes to overcome steric constraints.

### In-cell cryo-electron tomography confirms dominant co-co assembly in *trans* of PheA

To validate these observations in an unperturbed cellular context, we visualized PheA co-co assembly by cryo-electron tomography (cryo-ET) in *E. coli* (**Figure 4A**). WT *E. coli* or cells overexpressing either PheA or DDD were vitrified, thinned by cryo-focused ion beam (FIB) milling, and imaged by cryo-ET. Ribosomes were resolved up to ∼7.0 Å resolution by subtomogram averaging (**Figure 4B**, **S15**, **Supplementary Table 2**).

**Figure 4:**
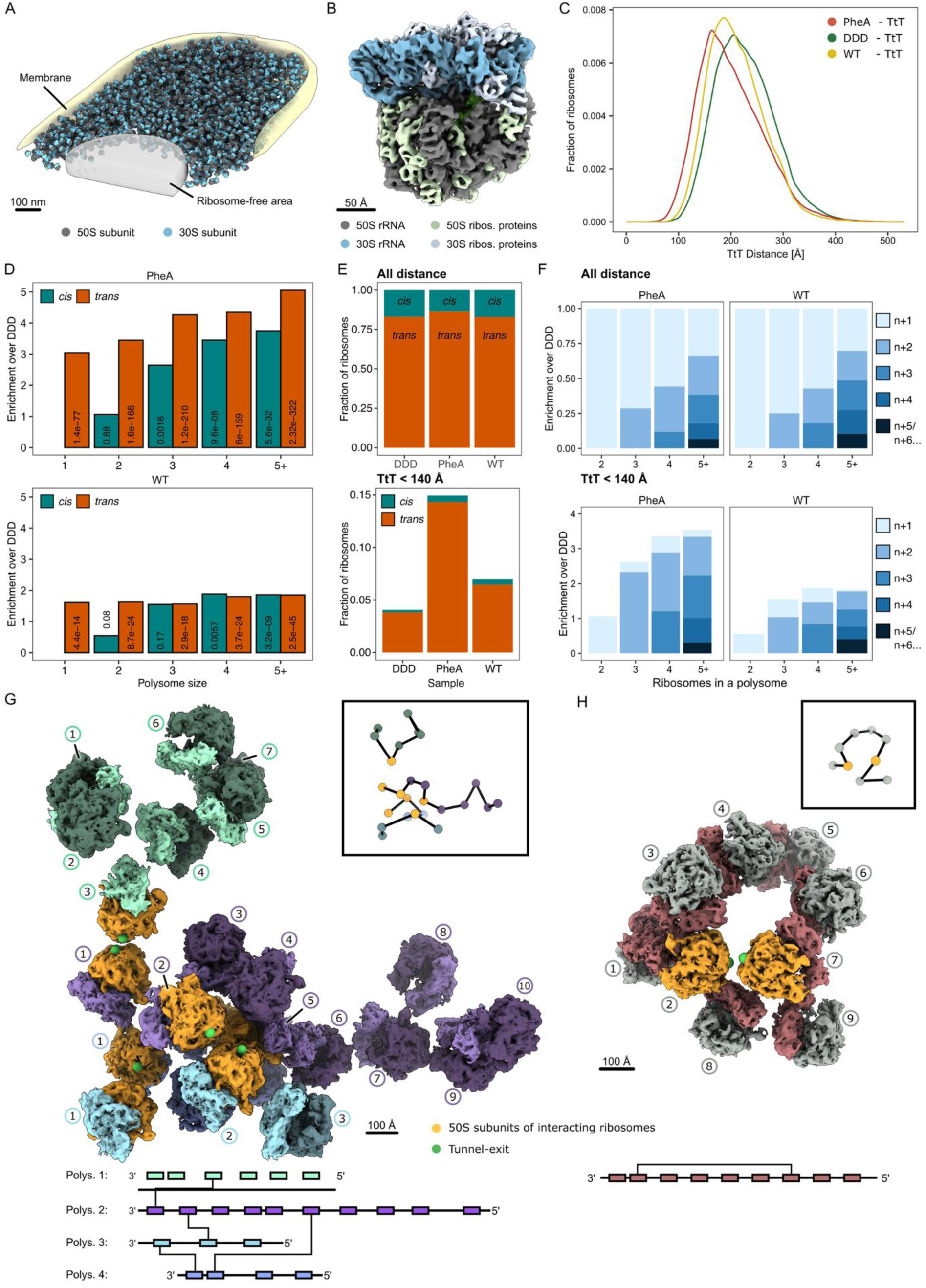
*Trans*-polypeptide exit tunnel proximity during co-co assembly is preserved *in situ*: A) 3D visualization of 70S ribosomes in the context of an *E. coli* cell (PheA). B) Cryo-ET reconstruction of 70S ribosomes at 7.4 Å resolution (PheA + DDD merged). C) Overall TtT distributions in PheA-(red), DDD-overexpressing (green) and WT cells (yellow). D) *In situ* enrichments *trans-* and *cis-*interactions of the PheA and WT sample over DDD within co-co distance, analyzed by the assigned polysome size. Fisher’s exact test p-values are indicated within each bar. E) Assignment of closest TtT neighbours on either same mRNA (*cis*) or different mRNA (*trans*) across all distances (above) or co-co distances (below). F) Enrichments of *cis*-interaction in PheA or WT samples over DDD across all distances (above) or co-co distances (below), analyzed by the assigned polysome size. The distribution of relative positions between closest neighbours (interaction partners) is indicated by the color within each bar. G) 3D visualization of four *trans*-interacting polysomes. Individual polysomes are coloured differently. H) 3D visualization of a *cis*-interacting polysome. Both G and H: The 50S subunits of interacting ribosomes are highlighted in orange; the tunnel exits by green spheres. The center-to-center traces of polysomes are shown in boxes. TtT interactions between polysomes are schematically illustrated below the 3D visualization.

Consistent with our cryo-EM analysis of isolated polysomes *in vitro* (**Figure 3B**), PheA-expressing cells showed a marked shift towards short TtT distances compared to WT and DDD-overexpressing cells (**Figure 4C**), confirming that polypeptide tunnel exit proximity drives co-co assembly *in vivo*. Compared to isolated polysomes, TtT distances in cells were overall shifted towards shorter distances, likely caused by the substantially higher local ribosome density in cells (**Figure S16A**,**B**). Notably, local ribosome density also slightly differed between the three tested conditions (**Figure S16C**). Therefore, to confirm that shorter TtT distances in PheA-overexpressing cells were a genuine effect of co-co assembly, we grouped ribosomes from all conditions into tiers with comparable local ribosome densities, and found that TtT distances within each tier remained significantly shorter in PheA-overexpressing cells (**Figure S16C**,**D**).

Mapping *cis* versus *trans* contacts in cells revealed that short TtT proximities occurred predominantly in *trans*, mirroring our *ex vivo* polysome data (**Figure 4D**,**E**). Notably, due to the high ribosome density in intact cells, we tested a range of mRNA entry-to-exit distance cut-offs for polysome assignment, observing robust *cis*/*trans* ratios for polysomal ribosomes (**Figure S17**). As before, short *cis* proximities depended strongly on high polysome occupancy and often linked ribosomes that were distant along the same mRNA (**Figure 4D**,**F**). Similar to our cryo-EM SPA data on isolated polysomes, higher enrichment of short TtT distances in PheA and WT over DDD samples was observed regardless of the TtT distance cut-off (**Figure S18**).

Finally, PheA polysomes in cells were visualized forming large, *trans*-connected ribosome networks (**Figure 4G**, **S19**), alongside rarer looping polysomes supporting *cis* contacts via distal ribosomes (**Figure 4H**; **S20**).

Together, visualizing these processes in cells confirms that PheA predominantly assembles co-translationally in *trans*, leveraging ribosome polypeptide exit tunnel proximity to form extensive polysome networks and demonstrating that *trans* co-co assembly can be a mechanism for co-translational complex formation of homomeric proteins.

## Discussion

Co-translational assembly is increasingly recognized as a key strategy to ensure efficient and accurate formation of multi-subunit protein complexes in cells. Previous work has highlighted *cis* co-co assembly as a mechanism for enforcing allele-specific oligomerization and limiting misfolding by coupling nascent chain synthesis to immediate subunit interaction (*11*). Here, by combining ribosome profiling, polysome fractionation, and cryo-EM structural analysis, we demonstrate that the bacterial homodimeric enzyme PheA can assemble in *trans*, that is, via oligomerization of nascent chains synthesized on different mRNAs. Using a dual-pull down assay we further show that PheA dimers can form from the translation products of two separate mRNA transcripts. These findings reveal that some homodimers can co-co assemble in *trans,* leading to the formation of interconnected mRNA networks in bacteria. Notably, PheA employs this mode particularly strongly, raising the question of what features might favor *trans* over *cis* interactions.

Our data suggest that PheA’s structural features create a unique dependency on rapid, correctly timed assembly. The PheA dimerization domain consists of three intertwined α-helices that are at the immediate N-terminus of the protein (**Figure 5A**). The intertwining structure of the two dimerization domains strongly suggests that folding depends critically on accurately timed inter-chain contacts. Failure to assemble promptly risks misfolding and aggregation, consistent with our observation that full-length PheA tends to form inclusion bodies during overexpression, whereas the dimerization-deficient mutant (DDD) retains higher solubility (**Figure S21**). According to our disome profiling analysis, PheA’s co-co assembly commences immediately after the dimerization domain emerges from the ribosomal tunnel exit – corresponding to codon #110 in the P-site of the translating ribosome - and is completed within the further translation of just 10 codons. This highlights an exceptionally narrow window for productive folding and assembly.

**Figure 5:**
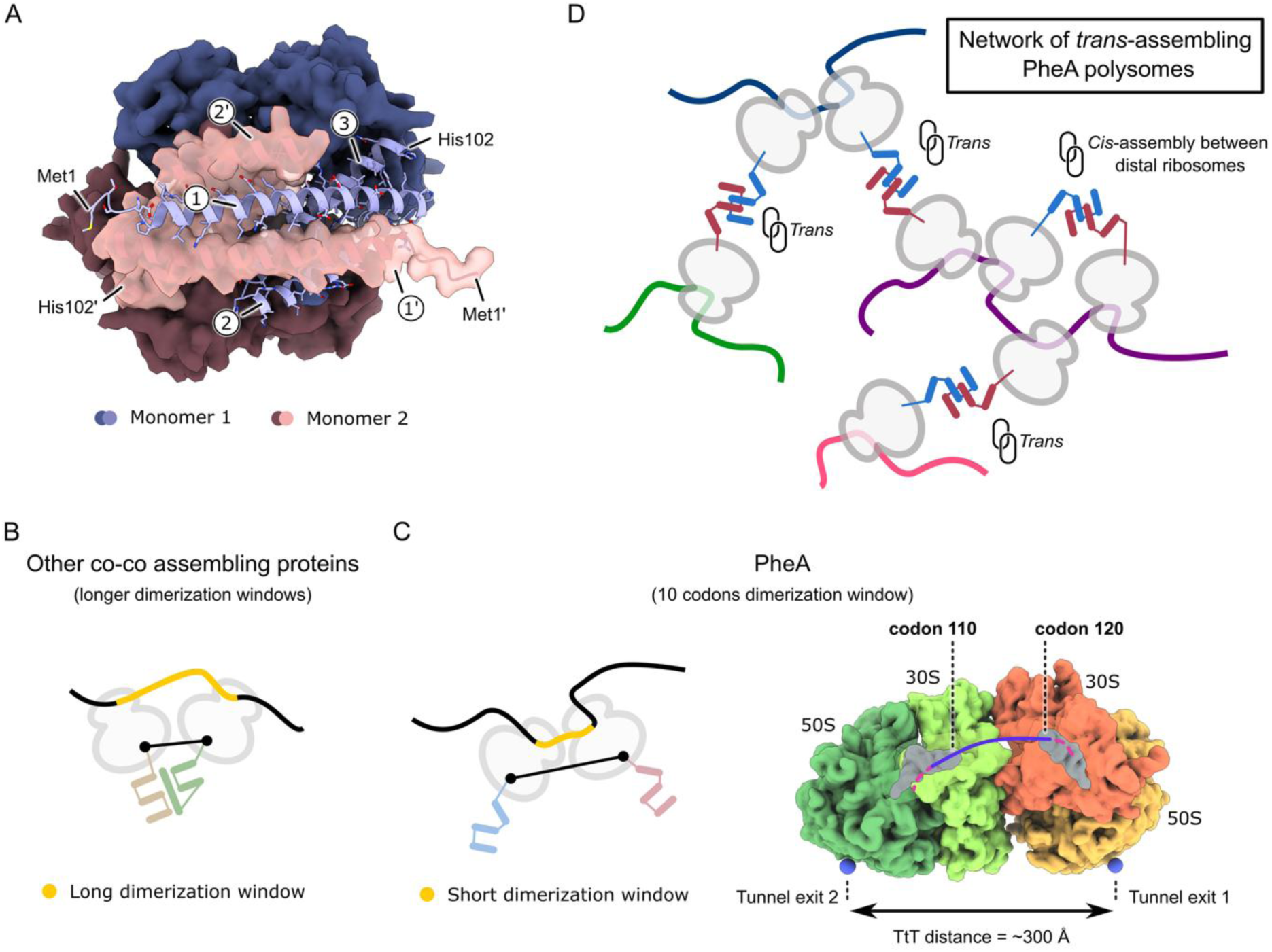
*Trans*-assembly can support rapid PheA complex formation in polysomes. A) Structure of the dimerized *E. coli* PheA dimerization domains (ranging from residues 1 – 89; residues 1 – 102 shown) (Alphafold 2 prediction). For dimerization domain of monomer 1 the surface is shown. First three alpha-helices forming the dimerization domain are indicated: Monomer 1: 1-3; Monomer 2: 1’-3’. B) Scheme illustrating how the simultaneous translation within the long dimerization windows of other co-co assembling proteins could allow for *cis* contacts. C) Left: Scheme illustrating how the simultaneous translation within the short dimerization windows of PheA orients the peptide exit tunnels of the involved ribosomes away from each other. Right: Simulated density representing the *B. subtilis* collided disome, in which the P-sites of leading and trailing ribosome are separated by ∼10 codons, corresponding to the PheA dimerization window (PDB: 7QV1 and 7QV2) (*48*). The 3D arrangement of such polysomal ribosomes is incompatible with *cis* contacts. D) Assembly in *trans* can form a network of interconnected polysomes*. Trans-*assembly can also support dimerization of nascent chains from lowly translated mRNA while higher packing or ribosomes on mRNA supports *cis-*assembly between distal ribosome.

This is in contrast to co-co assembling proteins with longer dimerization windows, which may allow the formation of kinked or other polysome topologies and thereby predominantly assemble via *cis* contacts of neighbouring ribosomes (**Figure 5B**). In contrast, for an envisioned *cis* assembly of PheA, the narrow dimerization window of PheA would place the mRNA exit of the first ribosome in very close proximity to the mRNA entry of the second ribosome (*31*, *32*) (**Figure 5C**), which ultimately limits the inter-ribosome flexibility below what is needed to accommodate alternative polysome topologies, like kinked or spiral organizations that may allow for polypeptide exit tunnel proximity.

We propose that PheA solves this topological constraint by predominantly assembling in *trans* via intertwining of the N-terminal helices, linking multiple polysomes into large, cross-connected ribosomal networks to increase the density of dimerization competent nascent chains (**Figure 5D**). To form co-co *trans* assembly disomes it may be sufficient to allow for free nascent subunit interactions, but we cannot exclude role of other, potentially proteinase K sensitive, factors driving these connected RNCs into the sucrose gradient pellet, such as chaperones or other nascent chain and ribosome binding factors bridging polysomes.

Our findings with PheA raise the broader question of how common *trans* co-co assembly might be in bacteria and eukaryotes. We identified four additional *E. coli* genes encoding homooligomers (AldA, ManX, GlpK, and AsnB) as candidates for forming *trans* networks (**Figure S3**). They all exhibit rapid assembly onsets during translation (<50 codons translated from basal level to maximum enrichment) and ORFs with substantial length (150–300 codons) downstream of the co-co assembly onset, features that, like PheA, may favor extensive and persistent *trans* interactions.

We also observed an enrichment of *trans* co-co ribosomes in the cryo-ET analysis of WT *E. coli* cells, but at a significantly lower level compared to PheA overexpression. Importantly, our results therefore do not argue that *trans* co-co assembly is the dominant pathway for homomers in general, but rather constitutes one co-co assembly mechanism in which small N-terminally located, intertwined dimerization domains of proteins drive co-co assembly towards *trans* co-co assembly to ensure rapid and productive dimerization. Thus, while the high ribosome density in the bacterial cytosol and the overall polysome architecture provide a suitable framework for *trans* assembly when allowed, we could only identify a small number of co-co assembling candidate proteins forming *trans*-networks in the entire *E. coli* proteome. This supports that *trans* co-co assembly is a specialized rather than a proteome-wide default pathway in *E. coli*.

While PheA appears to be a striking bacterial example, the underlying principles may be even more relevant in eukaryotes, where average ORF lengths are longer, facilitating potential multivalency of polysomes to form *trans*-networks. There is evidence for mRNA colocalization also outside of stress granules (*33*, *34*) and in the context of co-translational assembly (*6*, *7*, *35*), consistent with the possibility that distinct mRNA species can come into proximity to form co-co nascent chain interactions in *trans*.

Finally, our structural data reveal that the more rare *cis* co-co assembly of PheA occurs between distant polysomal neighbours under overexpression conditions – possibly as a consequence of local competition between nascent chains from the same and co-localized mRNA transcripts. The rare cases of *cis* co-co assembly, however, agree with our current knowledge of polysome topology, in which densely packed polysomes show polypeptide exit tunnels facing away from each other (*24*, *25*, *28*) (**Figure 5C**), while looser polysome arrangements may allow for close appositions of ribosomal peptide tunnel exits.

Such a polysomal organization is expected to limit opportunities for non-native interactions between nascent chains in densely packed polysomes while still permitting *cis* co-co assembly when spatial and translational constraints allow. In contrast, *trans* assembly provides an elegant solution for proteins such as PheA, whose interaction domain must assemble within an exceptionally narrow translational window and whose folding appears to depend on rapid intermolecular association. For the vast majority of proteins, however, such stringent spatial and temporal constraints are unlikely to exist, reducing the selective pressure to promote *trans* interactions. Instead, restricting assembly to less densely packed ribosomes translating the same mRNA may improve assembly fidelity by promoting *cis* assembly minimizing opportunities for promiscuous interactions with nascent chains synthesized elsewhere in the cell (*5*, *11*).

Together, these findings expand our understanding of how translation, mRNA localization, and ribosome organization collectively shape the timing and fidelity of protein complex assembly in cells. Beyond PheA, *trans* co-co assembly may represent a broader, underappreciated mechanism that cells use to coordinate the formation of homomeric complexes that must assemble quickly to avoid misfolding. Future work will define how widespread such ribosome– ribosome networks are, and how cells may tune mRNA abundance, supramolecular ribosome organization, and local translation environments to favor either *cis* or *trans* assembly pathways as needed.

## Materials and methods

### Used strains and plasmids

We have used *E. coli* strain MC4100 whenever wild-type cells (WT) are mentioned and BL21 (DE3) strains for overexpression experiments. For cloning and plasmid maintenance, we used *E. coli* XL1 Blue.

The plasmid for overexpression of PheA and its variants was cloned using Gibson assembly of pET3 vector and *pheA* gene amplified from wild-type *E. coli* genomic DNA. Non-dimerizing variants of PheA were cloned using either site-directed mutagenesis (PheA LLALL21A5) or circularization of linear PCR product via Gibson assembly (DDD variant, PheA_96-386_). The correctness of all the cloning was assessed by Sanger sequencing using T7 promotor primers.

### Cell growth and harvest

Cells were grown on Luria Broth (LB) plates and in liquid LB medium at 37 °C with the addition of 100 ug/mL of AMP. Liquid cultures were grown with orbital shaking at 130 rpm until an optical density (O.D.) of 0.3-0.5. Overexpression was induced by the addition of Isopropyl β-D-1-thiogalactopyranoside (IPTG) to the final concentration of 1 mM. After 15 minutes, cells were harvested by rapid filtration, scraped with a spatula and plunged in liquid nitrogen. For mild proteinase K digestion of MNase-resistant disomes, cells were harvested by centrifugation (4000 x g, 10 minutes) in the presence of 1 mM chloramphenicol (CAM). Pelleted cells were scraped with a spatula and plunged in liquid nitrogen.

WT cells were grown without antibiotics and directly harvested when O.D. reached 0.3-0.5.

### Cell lysis

Lysis buffer (50 mM HEPES.KOH, pH = 7.5, 150 mM KCl, 10 mM MgCl_2_, 5 mM CaCl_2_, 0.1 % Tween 20, 0.4 % NP-40 substitute, cOmplete protease inhibitors and DNase and 1 mM CAM) was frozen into small pellets and 500 uL were added into scraped cells on liquid nitrogen. Cells were lysed in precooled cryo-mill at 30 Hz for 2 minutes and the resulting powder was stored at -80 °C until further use.

### Polysome profiling and sequencing of sucrose gradient pellet fraction

Cell powder (n = 4) was quickly thawed in a water bath and by addition of 500 uL of lysis buffer. The lysate was clarified by centrifugation at 20 000 x g for 3 minutes. Clarified lysate was then treated either with proteinase K (1 mg of proteinase K per 200 mg of nucleic acid material) or buffer for 10 minutes on ice. The digestion was stopped by addition of 1 mM PMSF. The lysate was overlayed on a 15-60 % sucrose gradient in (50 mM HEPES.KOH, pH = 7.5, 150 mM KCl, 5 mM MgCl_2_, 0.1 mM CAM) and resolved by spinning at 35000 rpm in a SW-40 rotor at 4 °C for 3.5 hours. The gradient was fractionated on a Piston Fractionator (BioComp Instruments, USA) into 20 equivolume fractions. The remaining solution (roughly 700 uL) on the bottom of the tube was pipetted several times in the tube to resuspend a potential pellet in the sucrose gradient. CaCl_2_ was added to a final concentration of 5 mM and mRNA was digested by MNase (150 U per 1 A.U. at 260 nm) for 6 minutes at 25 °C. The reaction was quenched by adding 6 mM EGTA and the solution was mixed with equal volume of resuspension buffer (50 mM HEPES, 150 mM KCl, 5 mM MgCl2, pH = 7.5) to bring down the final sucrose concentration to roughly 30 %. Ribosomes pelleted by centrifugation at 100 000 rpm in SAT2 rotor for 1 hour were resuspended in 100 uL resuspension buffer and RNA was extracted by trizol reagent. Sequencing libraries were prepared as described (*36*).

### Plasmid Construction

To investigate trans-assembly of PheA, a plasmid named pETDuet_pheA containing two independently T7 promoter-expressed *pheA* ORFs was constructed. One PheA contains a C-terminal Strep-Tag, while the second PheA contains a C-terminal *trxA* and 3xFlag tag. The previously described pET-Duet-1 plasmid was used as the parental vector (*37*). The original ORF sequences were replaced with *pheA* sequences using two synthetic DNA fragments ordered from Twist Bioscience (USA). The fragments contained 20-nt overlaps to facilitate assembly. The vector and two *pheA*-containing fragments were assembled by three-fragment Gibson assembly using the Gibson Assembly Master Mix (NEB). Following assembly, the reaction was transformed into *E. coli* and plated on LB agar containing 100 µg/mL AMP. Individual colonies were isolated following overnight incubation at 37°C, and plasmid DNA was prepared and verified by Oxford Nanopore Technologies sequencing (ONT).

### Protein expression and Cell lysis

Plasmid pETDuet_pheA was freshly transformed into 50 µL of BL21 (DE3) competent cells. Following transformation, cells were plated on LB agar containing 100 µg/mL AMP and incubated overnight at 37°C. Individual colonies were inoculated into 5 mL LB supplemented with 100 µg/mL AMP and grown overnight at 37°C with shaking.

The following day, 200 mL LB cultures supplemented with 100 µg/mL AMP were inoculated from the overnight starter cultures to an initial OD600 of 0.025. Cultures were grown until an OD600 of approximately 0.12, at which point expression was induced with 0, 0.05, or 1 mM IPTG. Cultures were subsequently grown until the uninduced control reached an OD600 of 0.8, at which point cells from all conditions were harvested by centrifugation (10 min, 4,500 × g at 4°C, FiberLite F14-6 × 250y, 5,400 rpm).

Cell pellets were resuspended in lysis buffer (50 mM HEPES pH7.0, 150 mM KCl, 10 mM MgCl2, 5mM CaCl2, 0.4% Triton X-100, 0.1% NP-40, 1x cOmplete protease inhibitor). at a concentration corresponding to 50OD600 per mL. The cell suspensions were aliquoted into 1-mL portions and frozen in metal mixer-mill jars. Frozen cells were disrupted using a mixer mill (1 × 2 min, 30 Hz). Samples were either stored at −70°C or processed immediately for affinity purification.

Frozen samples were thawed at 25°C. MNase was added to a final amount of 2500 U, and samples were incubated for 5 min on ice. An aliquot of the lysate was collected prior to clarification. Lysates were clarified by centrifugation 20,000 × g for 10 min to remove insoluble material and aggregates. The resulting supernatants were collected as clarified lysates for affinity purification.

### Flag and Strep Affinity Purification

For affinity purification, 50 µL of either Strep-Tactin 4Flow beads (IBA) or ANTI-FLAG® M2 Affinity Gel beads (Millipore) were used per sample. Beads were first washed with 1 mL of binding buffer (50 mM HEPES-KOH, pH 7.0, 150 mM KCl). Clarified lysates were added to the beads and incubated for 30 min at 6°C with rotation at 30 rpm using an F1 overhead roller. Following incubation, beads were collected by centrifugation at 1,000 × g for 1 min, and the supernatant was retained as the unbound fraction.

Beads were washed with 1 mL of wash buffer (50 mM HEPES pH 7.0, 500 mM KCl, 0.4% triton X-100, 0.1% NP-40). and transferred to a fresh tube. Samples were incubated for 5 min at 4°C with rotation at 30 rpm. The washing procedure was repeated four additional times, for a total of five washes.

FLAG-bound proteins were eluted by incubation with 100 µg/mL FLAG peptide (ChromoTek 3xDYKDDDDK-peptide) in 50 mM HEPES-KOH, pH 7.0, 150 mM KCl. Strep-tagged proteins were eluted using 1× Strep-Tactin XT Elution Buffer BXT (IBA). Samples were incubated for 5 min at 25°C and subsequently centrifuged at 1,000 × g for 1 min. Approximately 80 µL of the resulting supernatant was collected as the elution fraction and immediately added to SDS – sample buffer.

### Western blotting

Samples were resolved on 4–12% SurePAGE SDS-PAGE gel (Genscript, M00653) in MES running buffer (Genscript, M00677). Immediately following electrophoresis, proteins were transferred to a PVDF membrane using the Trans-Blot Turbo transfer system (Bio-Rad, 1704150EDU) with Trans-Blot Turbo Mini PVDF transfer packs (Bio-Rad, 1704156). Transfer was performed for 30 min at 25 V and 1 A.

The membrane was blocked for 1 h in 3% BSA (Sigma-Aldrich, A7030) prepared in TBS-T and subsequently incubated overnight with anti-FLAG primary antibody (rabbit polyclonal anti-DYKDDDDK tag antibody, Proteintech, 20543-1-AP; 1:1,000 dilution).

The following day, the membrane was washed three times with TBS-T (50 mM Tris-HCl, pH 7.5, 150 mM NaCl, 0.1% Tween-20) for 10–30 min per wash. The membrane was then incubated for 10 min in biotin-blocking buffer (2.5 mg/mL avidin, 0.1% sodium azide), followed by a 1 h incubation with anti-rabbit AP-conjugated secondary antibody (Vector, AP-1000; 1:20,000 dilution) and AP-conjugated streptavidin (IBA, 2-1503-001; 1:4,000 dilution), allowing to visualize both PheA variants. The membrane was subsequently washed three times with TBS-T for 10–30 min per wash. ECF substrate (GE Healthcare Life Sciences, RPN3685) was applied to the membrane, and the blot was imaged after 5 min using an Amersham ImageQuant 600 imaging system (GE Healthcare Life Sciences, AI600, 29-0834-61).

### Isolating monosomes and disomes

Cell powder was quickly thawed in a water bath and by addition of 500 uL of lysis buffer. The thawed lysate was incubated with MNase (final concentration 150 U per 1 A.U. of nucleic acid material) and incubated at 25 °C for 12 minutes. The reaction was quenched by addition of 6 mM EGTA and clarified by centrifugation at 4 °C for 3 minutes at 20000 x g. The clarified lysate was either further processed (see below) or layered on top of a 15-45 % sucrose gradient (50 mM HEPES.KOH, pH = 7.5, 150 mM KCl, 5 mM MgCl_2_, 0.1 mM CAM) and resolved by spinning in a SW-40 rotor at 35 000 rpm for 3.5 hours. The gradient was fractionated on Piston Fractionator (BioComp Instruments, USA), fractions containing monosomes and disomes were further processed according to further application (see below).

### Mild proteinase K digestion of MNase-treated lysates

Nuclease digested lysate was incubated with proteinase K solution (prepared from powder, dissolved in lysis buffer to an approximate concentration of 10 mg/mL). Several dilutions of proteinase K were added to the lysate so that the final dilution of proteinase K reached 1:200, 1:600, 1:1800, or 1:5400 (mg of protein per mg of nucleic acid material). The digestion was performed on ice for 10 minutes, after which 1 mM PMSF (final) was added to the lysate. The reaction was clarified by centrifugation at 20 000 x g for 3 minutes and layered on top of 5-45 % sucrose gradient and processed as described above.

### Polysome profiling of PheA/DDD overexpressing cells

Cell powder of PheA/DDD overexpressing cells was quickly thawed in a water bath and by addition of 500 uL of lysis buffer. The thawed lysate was incubated with either lysis buffer or proteinase K (1 mg of protein per 200 mg of nucleic acid material), incubated on ice for 10 minutes and then quenched by 1 mM PMSF (final concentration). The lysate was clarified by centrifugation at 20 000 × g for 3 minutes. Clarified lysate was loaded on 15 – 60 % sucrose gradient (50 mM HEPES.KOH, pH = 7.5, 150 mM KCl, 5 mM MgCl_2_, 0.1 mM CAM) and resolved by spinning at 35000 rpm in a SW-40 rotor at 4 °C for 3.5 hours. The gradient was fractionated on a Piston Fractionator (BioComp Instruments, USA) into 20 equivolume fractions.

To test how clarification steps impact the ribosomal content of the PheA-overexpression lysate, we have divided thawed lysate into half. One was treated directly with MNase (150 U per 40 µg of RNA), quenched by 6 mM EGTA and then clarified by centrifugation at 20 000 × g for 3 minutes. The other half was clarified by centrifugation at 20 000 × g for 3 minutes, then treated with MNase (150 U per 40 µg of RNA) and quenched by 6 mM EGTA. Clarified lysates were loaded on 15 – 60 % sucrose gradient (50 mM HEPES.KOH, pH = 7.5, 150 mM KCl, 5 mM MgCl_2_, 0.1 mM CAM) and resolved by spinning at 35000 rpm in a SW-40 rotor at 4 °C for 3.5 hours. The gradients were fractionated on a Piston Fractionator (BioComp Instruments, USA) into 20 equivolume fractions.

### Disome selective profiling

Sucrose gradient fractions containing monosomes and disomes were pooled and extracted by phenol chlorophorm. Sequencing libraries (n = 2) were prepared and sequenced as described previously (*38*).

### DiSP analysis

3’ adaptor-trimmed reads were aligned to the MC4100 genome (build GCA_000499485.1). PCR duplicates were removed by utilizing 10N unique molecular identifiers (UMIs) using custom scripts as described before (*36*, *38*). Aligned data processing and visualization was done in R with Riboseqtools (*11*).

### Chorismate mutase assay

To test chorismate mutase activity, reactions containing 5 mM chorismate, 100 mM Tris, 10 mM MgCl_2_, 1 mM DTT, 0.5 mM EDTA, pH = 7.0 were mixed with either 40 ng/uL PheA disomes (originated from PheA overexpression), 40 ng/uL ribosomes (NEB), or water and incubated at 37 °C. After 0, 15, 32, and 110 minutes, UV absorption spectra were recorded on Nanodrop. The decrease in absorbance at 270 and 290 nm were evaluated as a hallmark of chorismate isomerization to prephenate (*39*).

### Sample preparation for cryo-EM SPA

For cryo-EM SPA of MNase resistant disomes, disome and monosome fractions used for DiSP were pooled and spined at 400000 x g for 90 minutes. After discarding the supernatant, the pellets were dissolved to a final concentration of 100 ng / uL of RNA (corresponding to roughly 50 nM ribosomal concentration).

For cryo-EM SPA of intact polysomes rapidly harvested frozen cells were pulverized in a cryomill in a lysis buffer without detergents and CAM (50 mM HEPES.KOH, pH = 7.5, 150 mM KCl, 10 mM MgCl_2_, 5 mM CaCl_2_ cOmplete protease inhibitors and DNase). Lysate powder was stored in -80 °C until use. Before vitrification, the lysate was quickly thawed in hand and by adding variable amount of lysis buffer to achieve sufficient dilution. Cell debris was removed by centrifugation at 20 000 x g for 3 minutes at 4 °C prior to vitrification.

### Cryo-EM grid preparation and data acquisition for SPA

For cryo-EM grid preparation, holey carbon grids (R2/1, copper, 200 mesh, Quantifoil Micro Tools) were glow-discharged in a PELCO easiGlow. EM grids for SPA were prepared using a Vitrobot Mark IV (ThermoFisher Scientific) at 4 °C and 100 % humidity: 5 µl of cell lysate was applied per grid and blotting before plunge-freezing in liquid ethane was conducted using Whatman filter paper Nr. 1 at a blot force of 10. The blotting time was 5 seconds.

All datasets were acquired at a Titan Krios TEM (ThermoFisher Scientific) using EPU (ThermoFisher Scientific). Data were acquired at 300kV acceleration voltage using an energy-filtered K3 camera (Gatan) in dose fractionation mode at a pixel size of 1.07 Å/px. The defocus range was set to -0.75 µm to -1.75 µm. In each selected hole 4 micrograph movie stacks containing 30 frames each were acquired with similar cumulative dose (WT: 45.26 e^-^/Å^2^; PheA: 43.6 e^-^/Å^2^; DDD: 45.00 e^-^/Å^2^).

### Cryo-EM grid for FIB milling and tilt series acquisition

Similar as for SPA cryo-EM grid preparation, *E. coli* cells carrying different plasmids were induced for PheA overexpression by IPTG addition for 15 min. *E. coli* cells from all conditions, including *E. coli* WT cells were centrifuged (8000 xg, 1 min, RT) immediately before sample application to the grid.

For cryo-EM grid preparation, holey carbon grids (R2/1, copper, 200 mesh, Quantifoil Micro Tools) were glow-discharged in a PELCO easiGlow. EM grids for cryo-ET were prepared using a Vitrobot Mark IV (ThermoFisher Scientific) at 37°C and at a humidity of approx. 30%. To obtain a specimen topology suited for cryo-FIB milling, the cell pellet was not resuspended after centrifugation and approx. 1 µl of cell pellet and approx. 4 µl of supernatant were applied to the cryo-EM grids. The grids were then blotted from one side only using Whatman filter paper Nr. 1 from the back side of the grid and a Fluoropolymer sheet (Science Service) from the front.

Vitreous lamellae of *E. coli* cells were prepared in an Aquilos 2 dual-beam cryo-FIB-SEM microscope (ThermoFisher Scientific) operated by Maps Software (ThermoFisher Scientific) in combination with AutoTEM (ThermoFisher Scientific). The specimen was coated with organometallic platinum for 60 seconds before milling. Lamellae with a target thickness of 140 nm to 150 nm were prepared in 5 successive steps using a Gallium-ion beam (acceleration voltage 30 kV) with step-wise decreasing current.

Cryo-ET data were acquired at a Titan Krios TEM (ThermoFisher Scientific) using Tomography 5 (ThermoFisher Scientific). The data were recorded at 300 kV acceleration voltage using an energy-filtered K3 camera (Gatan) in dose fractionation mode at a pixel size of 2.66 Å/px. Tilt series were acquired using a dose-symmetric tilt-scheme with a 3° increment starting from 7°, corresponding approximately to the pre-tilt of the milled lamellas. Each tilt image was fractionated into 5 frames with a per tilt exposure dose of 3.78 e^-^/Å^2^ for PheA, 3.9 e^-^/Å^2^ for DDD and 3.8 e^-^/Å^2^ for WT.

### Cryo-EM SPA image processing

Image processing was completely performed in Relion 3.1 (*40*) and is summarized in **Figure S7** and **Supplementary Table 1**. Micrograph movie stacks from each data set were separately subjected to motion correction using MotionCor2 (*41*) using a 5 x 5 grid and gCTF (*42*) to estimate the contrast transfer function. Micrographs from PheA were at this point additionally split into two subsets depending on the estimated maximal resolution (< 3.5 Å and 3.5 Å – 4.5 Å). For WT and PheA DDD only micrographs with an estimated maximal resolution below 3.5 Å were retained. Autopicking was performed on each micrograph subset and particles were extracted with a pixel size of 4.28 Å/px (128 px^3^). To sort for true-positive ribosomal particles, particles from the subsets DDD and PheA were subjected to 3D classifications using a 70S mask, while particles of the PheA subsets were subjected to three subsequent 3D classification runs with a spherical mask. After duplicate removal within a radius of 100 Å, true-positive particles from all three datasets were merged and subjected to 3D auto-refinement followed by a 3D classification without images alignment for rejection of 50S ribosomal subunits, both using a 70S mask.

### Subtomogram analysis

Subtomogram analysis was performed using Relion 3.1 (*40*) in combination with Warp/M (*43*, *44*) and in house-scripts. The subtomogram analysis workflow is summarized in **Figure S9** and **Supplementary Table 2**. Initial data processing for each dataset was conducted separately: Micrograph movie stacks were subjected to motion correction using MotionCor2 (*41*) using a 5 x 5 grid and pre-processing in Warp. Tilt series were aligned using patch-tracking in IMOD 4.11.25 (*45*). The obtained tilt series alignment parameters were imported into Warp for reconstruction of binned tomograms (21.28 Å/px). Template matching against 70S ribosomes was conducted using PyTom 1.1 (*46*) using a 70S ribosome as a template. 10 000 candidate positions were identified per tomogram and particle positions outside of the cells were removed manually using USCF ChimeraX (*47*). From the remaining positions, the top-scoring 40% in terms of cross-correlation values were retained. Subtomograms were reconstructed in Warp (128 px^3^, 4.0 Å/px) and aligned using 3D auto-refinement as implemented in Relion 3.1. Based on the refined particle positions and orientations, local tilt-series alignment was optimized in M (*44*). Using optimized tomograms, particle localization via template matching, candidate extraction and manual removal of candidate positions outside of cellular sections were repeated. For the remaining candidate positions, optimized subtomograms were reconstructed (128 px^3^, 4.0 Å/px) and subjected to 3D auto-refinement, followed by 3D classification without image alignment. Low-quality ribosome classes were re-classified using 3D classification with image alignment. The final subsets of true-positive 70S particles from all datasets were merged and subjected to 3D auto-refinement.

### Analysis of ribosome configurations

Ribosome configurations in cryo-EM SPA data and cryo-ET data were analyzed using the same workflow. For cryo-EM SPA data, the Z-position of particles was in a first step computed based on the estimated defocus. To compare measured distances from both cryo-EM modalities the atomic model of a *B. subtilis* 70S ribosomes (7QV1) (*31*) was fitted into the final cryo-EM reconstructions (**Figure S7**) and marker pairs were positioned either on the mRNA entry and exit or on the last residue of the nascent chain before leaving the tunnel-exit. For both SPA data and cryo-ET data, polysome tracing was performed as described in (*30*) based on mRNA entry and exit markers. TtT-distances were measured using the same approach but using markers on the tunnel exits of adjacent ribosomes. Local ribosome density was measured in shells of 40 nm, 100 nm, 300 nm, 500 nm and 1 µm.

### NND Cluster analysis

Nearest neighbour distance cluster analysis was performed using previously published scripts (*25*) in Matlab R2023a with the following modifications: Relion star file of MNase-digested disome sample was sub-setted for particles with their closest TtT neighbour within 140 Å and the first 32000 particles listed in the star file were used for analysis. As the sample is originated from SPA grids, the Z-coordinate was manually set to 0 for all particles. Equal weight was given to all parameters (i.e. distance, direction and orientation) during clustering and dendrogram distance cut-off was set at 2. The 10 most abundant clusters out of ∼600, accounting for ∼ 50% of analyzed particles, were visualized (see Fig. S6).

## Data availability

All cryo-EM reconstructions obtained in this study were deposited in the Electron Microscopy Data Bank (EMDB) with the following accession codes: EMD-55639 (monosome from MNase-digested polysomes), EMD-55640 (disome from MNase-digested poylsomes), EMD-55641 (consensus 70S ribosome from polysomes), EMD-55642 (consensus 70S ribosome from cellular cryo-ET (PheA + DDD)), EMD-58666 (consensus 70S ribosome from cellular cryo-ET (WT)). Custom scripts used in this study are available as Supplementary Material. Ribosome profiling data has been deposited to the GEO databank under the accession code GSE303296.

## Acknowledgements

We acknowledge the services SDS@hd and bwHPC supported by the Ministry of Science, Research and the Arts Baden-Württemberg, as well as projects INST 35/1314-1 FUGG and INST 35/1597-1 FUGG funded by the German Research Foundation (DFG). We acknowledge access to the infrastructure and support provided by the Cryo-EM Network at Heidelberg University (HDcryoNet), which is funded by the DFG, the Federal Ministry of Education and Research (BMBF) and the Ministry of Science Baden-Württemberg, among others, within the framework of the Excellence Strategy of the Federal and State Governments of Germany. This work is supported by grants of the German-Israeli Foundation for Scientific Research and Development (I-1564-417.13/2023) to G.K. and the European Research Council (ERC) to S.P. (ERC-StG - 101075851 - RiboStress) and to B.B. (ERC - SyG - 101072047 - CoTransComplex). Views and opinions expressed are, however, those of the authors only and do not necessarily reflect those of the European Union or the ERC. Neither the European Union nor the granting authority can be held responsible for them. S.P. acknowledges funding by the Aventis Foundation and the Chica and Heinz Schaller Foundation. J.K. was supported by Horizon 2020 MSCA Individual Fellowship (CoCoAssembly 895164). J.S. was supported by a PhD fellowship from the Studienstiftung des deutschen Volkes (German Academic Scholarship Foundation) and a completion grant from the Baden-Württemberg State Graduate Funding (LGF). J.S. is a member of the Heidelberg Biosciences International Graduate School (HBIGS).

## Author contributions

J.K. performed cell culture, polysome profiling, ribosome profiling libraries with the exception of MC4100 DiSP libraries which were prepared by J.S. G.F. performed the dual-pull down assay. S.F. and J.K. prepared cryo-EM specimens and S.F. collected all cryo-EM data. Cryo-ET data was collected, processed and analysed by S.F. Cryo-EM SPA data was processed by J.K. with assistance from S.F. Customs scripts were developed by S.F. and S.K.. S.P., G.K. and B.B. supervised the experiments. J.K., S.F. and S.P. wrote the manuscript with revisions from G.K. and B.B.. All authors discussed the data and gave final approval for publication.

## Reference

1. A. J. Reid, J. A. Ranea, C. A. Orengo, Comparative evolutionary analysis of protein complexes in E. coli and yeast. BMC Genomics 11, 79 (2010).

2. M. Lynch, The Evolution of Multimeric Protein Assemblages. Molecular Biology and Evolution 29, 1353–1366 (2012).

3. S. Kassem, Z. Villanyi, M. A. Collart, Not5-dependent co-translational assembly of Ada2 and Spt20 is essential for functional integrity of SAGA. Nucleic Acids Research 45, 1186–1199 (2017).

4. Y.-W. Shieh, P. Minguez, P. Bork, J. J. Auburger, D. L. Guilbride, G. Kramer, B. Bukau, Operon structure and cotranslational subunit association direct protein assembly in bacteria. Science 350, 678–680 (2015).

5. A. Shiber, K. Döring, U. Friedrich, K. Klann, D. Merker, M. Zedan, F. Tippmann, G. Kramer, B. Bukau, Cotranslational assembly of protein complexes in eukaryotes revealed by ribosome profiling. Nature 561, 268–272 (2018).

6. O. O. Panasenko, S. P. Somasekharan, Z. Villanyi, M. Zagatti, F. Bezrukov, R. Rashpa, J. Cornut, J. Iqbal, M. Longis, S. H. Carl, C. Peña, V. G. Panse, M. A. Collart, Co-translational assembly of proteasome subunits in NOT1-containing assemblysomes. Nat Struct Mol Biol 26, 110–120 (2019).

7. S. Mallik, J. Venezian, A. Lobov, M. Heidenreich, H. Garcia-Seisdedos, T. O. Yeates, A. Shiber, E. D. Levy, Structural determinants of co-translational protein complex assembly. Cell 188, 764–777.e22 (2025).

8. M. Seidel, A. Becker, F. Pereira, J. J. M. Landry, N. T. D. De Azevedo, C. M. Fusco, E. Kaindl, N. Romanov, J. Baumbach, J. D. Langer, E. M. Schuman, K. R. Patil, G. Hummer, V. Benes, M. Beck, Co-translational assembly orchestrates competing biogenesis pathways. Nat Commun 13, 1224 (2022).

9. S. Wagner, A. Herrmannová, V. Hronová, S. Gunišová, N. D. Sen, R. D. Hannan, A. G. Hinnebusch, N. E. Shirokikh, T. Preiss, L. S. Valášek, Selective Translation Complex Profiling Reveals Staged Initiation and Co-translational Assembly of Initiation Factor Complexes. Molecular Cell 79, 546–560.e7 (2020).

10. G. Yayli, A. Bernardini, P. K. Mendoza Sanchez, E. Scheer, M. Damilot, K. Essabri, B. Morlet, L. Negroni, S. D. Vincent, H. T. M. Timmers, L. Tora, ATAC and SAGA co-activator complexes utilize co-translational assembly, but their cellular localization properties and functions are distinct. Cell Reports 42, 113099 (2023).

11. M. Bertolini, K. Fenzl, I. Kats, F. Wruck, F. Tippmann, J. Schmitt, J. J. Auburger, S. Tans, B. Bukau, G. Kramer, Interactions between nascent proteins translated by adjacent ribosomes drive homomer assembly. Science 371, 57–64 (2021).

12. F. Wruck, J. Schmitt, K. Fenzl, M. Bertolini, A. Katranidis, B. Bukau, G. Kramer, S. Tans, Co-translational ribosome pairing enables native assembly of misfolding-prone subunits. Molecular Biology [Preprint] (2023). 10.1101/2023.06.30.547139.

13. C. M. Brennan, L. P. Vaites, J. N. Wells, S. Santaguida, J. A. Paulo, Z. Storchova, J. W. Harper, J. A. Marsh, A. Amon, Protein aggregation mediates stoichiometry of protein complexes in aneuploid cells. Genes Dev. 33, 1031–1047 (2019).

14. S. Juszkiewicz, R. S. Hegde, Quality Control of Orphaned Proteins. Molecular Cell 71, 443–457 (2018).

15. U. Schubert, L. C. Antón, J. Gibbs, C. C. Norbury, J. W. Yewdell, J. R. Bennink, Rapid degradation of a large fraction of newly synthesized proteins by proteasomes. Nature 404, 770–774 (2000).

16. A. Bernardini, L. Tora, Co-translational Assembly Pathways of Nuclear Multiprotein Complexes Involved in the Regulation of Gene Transcription. Journal of Molecular Biology 436, 168382 (2024).

17. A. Bernardini, P. Mukherjee, E. Scheer, I. Kamenova, S. Antonova, P. K. Mendoza Sanchez, G. Yayli, B. Morlet, H. T. M. Timmers, L. Tora, Hierarchical TAF1-dependent co-translational assembly of the basal transcription factor TFIID. Nat Struct Mol Biol 30, 1141– 1152 (2023).

18. G. Yayli, A. Bernardini, P. K. Mendoza Sanchez, E. Scheer, M. Damilot, K. Essabri, B. Morlet, L. Negroni, S. D. Vincent, H. T. M. Timmers, L. Tora, ATAC and SAGA co-activator complexes utilize co-translational assembly, but their cellular localization properties and functions are distinct. Cell Reports 42, 113099 (2023).

19. K. Till, V. Sunderlikova, F. Tippmann, P. Jevtić, M. Bertolini, K. Fenzl, J. Schmitt, A. Katranidis, B. Bukau, G. Kramer, M. Rapé, S. Tans, Translation-driven temporal control for intertwined protein assembly. Molecular Biology [Preprint] (2025). 10.1101/2025.08.25.672138.

20. F. Wruck, J. Schmitt, K. Till, K. Fenzl, M. Bertolini, F. Tippmann, A. Katranidis, B. Bukau, G. Kramer, S. J. Tans, Co-translational ribosome pairing enables native assembly of misfolding-prone subunits. Nat Commun 16, 7626 (2025).

21. A. Roeselová, S. Shivakumaraswamy, G. Jurkeviciute, J. Z. He, J. Auburger, J. L. Schmitt, G. Kramer, B. Bukau, R. I. Enchev, D. Balchin, The ribosome synchronizes folding and assembly to promote oligomeric protein biogenesis. Biochemistry [Preprint] (2025). 10.1101/2025.05.27.656346.

22. M. Metelev, E. Lundin, I. L. Volkov, A. H. Gynnå, J. Elf, M. Johansson, Direct measurements of mRNA translation kinetics in living cells. Nat Commun 13, 1852 (2022).

23. M. Badonyi, J. A. Marsh, Buffering of genetic dominance by allele-specific protein complex assembly. Sci. Adv. 9, eadf9845 (2023).

24. F. Brandt, S. A. Etchells, J. O. Ortiz, A. H. Elcock, F. U. Hartl, W. Baumeister, The Native 3D Organization of Bacterial Polysomes. Cell 136, 261–271 (2009).

25. F. Brandt, L.-A. Carlson, F. U. Hartl, W. Baumeister, K. Grünewald, The Three-Dimensional Organization of Polyribosomes in Intact Human Cells. Molecular Cell 39, 560– 569 (2010).

26. Z. A. Afonina, A. G. Myasnikov, V. A. Shirokov, B. P. Klaholz, A. S. Spirin, Conformation transitions of eukaryotic polyribosomes during multi-round translation. Nucleic Acids Research 43, 618–628 (2015).

27. A. G. Myasnikov, Z. A. Afonina, J.-F. Ménétret, V. A. Shirokov, A. S. Spirin, B. P. Klaholz, The molecular structure of the left-handed supra-molecular helix of eukaryotic polyribosomes. Nat Commun 5, 5294 (2014).

28. L. Xue, S. Lenz, M. Zimmermann-Kogadeeva, D. Tegunov, P. Cramer, P. Bork, J. Rappsilber, J. Mahamid, Visualizing translation dynamics at atomic detail inside a bacterial cell. Nature 610, 205–211 (2022).

29. A. Y. Lee, P. A. Karplus, B. Ganem, J. Clardy, Atomic structure of the buried catalytic pocket of Escherichia coli chorismate mutase. J. Am. Chem. Soc. 117, 3627–3628 (1995).

30. K. Helena-Bueno, S. Kopetschke, S. Filbeck, L. I. Chan, S. Birsan, A. Baslé, M. Hudson, S. Pfeffer, C. H. Hill, S. V. Melnikov, Structurally heterogeneous ribosomes cooperate in protein synthesis in bacterial cells. Nat Commun 16, 2751 (2025).

31. F. Cerullo, S. Filbeck, P. R. Patil, H.-C. Hung, H. Xu, J. Vornberger, F. W. Hofer, J. Schmitt, G. Kramer, B. Bukau, K. Hofmann, S. Pfeffer, C. A. P. Joazeiro, Bacterial ribosome collision sensing by a MutS DNA repair ATPase paralogue. Nature 603, 509–514 (2022).

32. K. Saito, H. Kratzat, A. Campbell, R. Buschauer, A. M. Burroughs, O. Berninghausen, L. Aravind, R. Green, R. Beckmann, A. R. Buskirk, Ribosome collisions induce mRNA cleavage and ribosome rescue in bacteria. Nature 603, 503–508 (2022).

33. X. Pichon, A. Bastide, A. Safieddine, R. Chouaib, A. Samacoits, E. Basyuk, M. Peter, F. Mueller, E. Bertrand, Visualization of single endogenous polysomes reveals the dynamics of translation in live human cells. Journal of Cell Biology 214, 769–781 (2016).

34. B. Wu, C. Eliscovich, Y. J. Yoon, R. H. Singer, Translation dynamics of single mRNAs in live cells and neurons. Science 352, 1430–1435 (2016).

35. I. Kamenova, P. Mukherjee, S. Conic, F. Mueller, F. El-Saafin, P. Bardot, J.-M. Garnier, D. Dembele, S. Capponi, H. T. M. Timmers, S. D. Vincent, L. Tora, Co-translational assembly of mammalian nuclear multisubunit complexes. Nat Commun 10 (2019).

36. J. Koubek, K. Jetzinger, S. Dror, M. Irastortza-Olaziregi, D. Frank, I. Kotan, J. Santos, F. Tippmann, P. Lafrenz, H. Kaessmann, O. Amster-Choder, B. Bukau, G. Kramer, A simple, fast and cost-efficient protocol for ultra-sensitive ribosome profiling. Cold Spring Harbor Laboratory [Preprint] (2025). 10.1101/2025.04.09.647970.

37. D. Mermans, F. Nicolaus, K. Fleisch, G. Von Heijne, Cotranslational folding and assembly of the dimeric *Escherichia coli* inner membrane protein EmrE. Proc. Natl. Acad. Sci. U.S.A. 119, e2205810119 (2022).

38. L. Eismann, I. Fijalkowski, C. V. Galmozzi, J. Koubek, F. Tippmann, P. Van Damme, G. Kramer, Selective ribosome profiling reveals a role for SecB in the co-translational inner membrane protein biogenesis. Cell Reports 41, 111776 (2022).

39. G. L. E. Koch, D. C. Shaw, F. Gibson, The purification and characterisation of chorismate mutase-prephenate dehydrogenase from Escherichia coli K12. Biochimica et Biophysica Acta (BBA) - Protein Structure 229, 795–804 (1971).

40. J. Zivanov, T. Nakane, S. H. W. Scheres, Estimation of high-order aberrations and anisotropic magnification from cryo-EM data sets in *RELION* -3.1. IUCrJ 7, 253–267 (2020).

41. S. Q. Zheng, E. Palovcak, J.-P. Armache, K. A. Verba, Y. Cheng, D. A. Agard, MotionCor2: anisotropic correction of beam-induced motion for improved cryo-electron microscopy. Nat Methods 14, 331–332 (2017).

42. K. Zhang, Gctf: Real-time CTF determination and correction. Journal of Structural Biology 193, 1–12 (2016).

43. D. Tegunov, P. Cramer, Real-time cryo-electron microscopy data preprocessing with Warp. Nat Methods 16, 1146–1152 (2019).

44. D. Tegunov, L. Xue, C. Dienemann, P. Cramer, J. Mahamid, Multi-particle cryo-EM refinement with M visualizes ribosome-antibiotic complex at 3.5 Å in cells. Nat Methods 18, 186–193 (2021).

45. J. R. Kremer, D. N. Mastronarde, J. R. McIntosh, Computer Visualization of Three-Dimensional Image Data Using IMOD. Journal of Structural Biology 116, 71–76 (1996).

46. M. L. Chaillet, G. van der Schot, I. Gubins, S. Roet, R. C. Veltkamp, F. Förster, Extensive Angular Sampling Enables the Sensitive Localization of Macromolecules in Electron Tomograms. Int J Mol Sci 24, 13375 (2023).

47. E. C. Meng, T. D. Goddard, E. F. Pettersen, G. S. Couch, Z. J. Pearson, J. H. Morris, T. E. Ferrin, UCSF CHIMERAX : Tools for structure building and analysis. Protein Science 32, e4792 (2023).

48. F. Cerullo, S. Filbeck, P. R. Patil, H.-C. Hung, H. Xu, J. Vornberger, F. W. Hofer, J. Schmitt, G. Kramer, B. Bukau, K. Hofmann, S. Pfeffer, C. A. P. Joazeiro, Bacterial ribosome collision sensing by a MutS DNA repair ATPase paralogue. Nature 603, 509–514 (2022).

