## Supplementary Figures and Tables for "Structural basis for co-translational assembly of homo-oligomeric proteins in *cis* and in *trans*"



**Abstract**

The efficient formation of protein complexes is essential for cellular function. Across all kingdoms of life, protein complexes frequently assemble co-translationally, either when one subunit is fully synthesized before interaction with nascent partner subunits (co-post) or when multiple nascent subunits interact co-translationally (co-co), with homomeric proteins particularly enriched among co-co assembling complexes. However, it is unknown whether co-co assembly of homomers occurs on the same mRNA in *cis* or across neighbouring mRNAs in *trans*, and how ribosomes spatially organize to enable co-co assembly. Using *E. coli* homodimeric protein PheA as a proof-of-concept model, we show by ribosome profiling and cryo-EM structural analysis that it employs the co-co assembly route. Co-co assembly of PheA is facilitated by the proximity of polypeptide exit tunnels, but does not rely on fixed ribosomal orientations. Surprisingly, PheA was identified as an exceptional candidate where *trans*-assembly is potentially the primary mode of PheA co-co assembly, generating large polysomal networks for PheA synthesis. *Cis*-assembly of PheA was less frequent and occurred mainly between non-adjacent ribosomes on the mRNA due to spatial restraints imposed by the arrangement of directly neighbouring ribosomes in polysomes. These findings reveal fundamental principles of how cells structurally organise fast and efficient co-translational protein complex assembly.

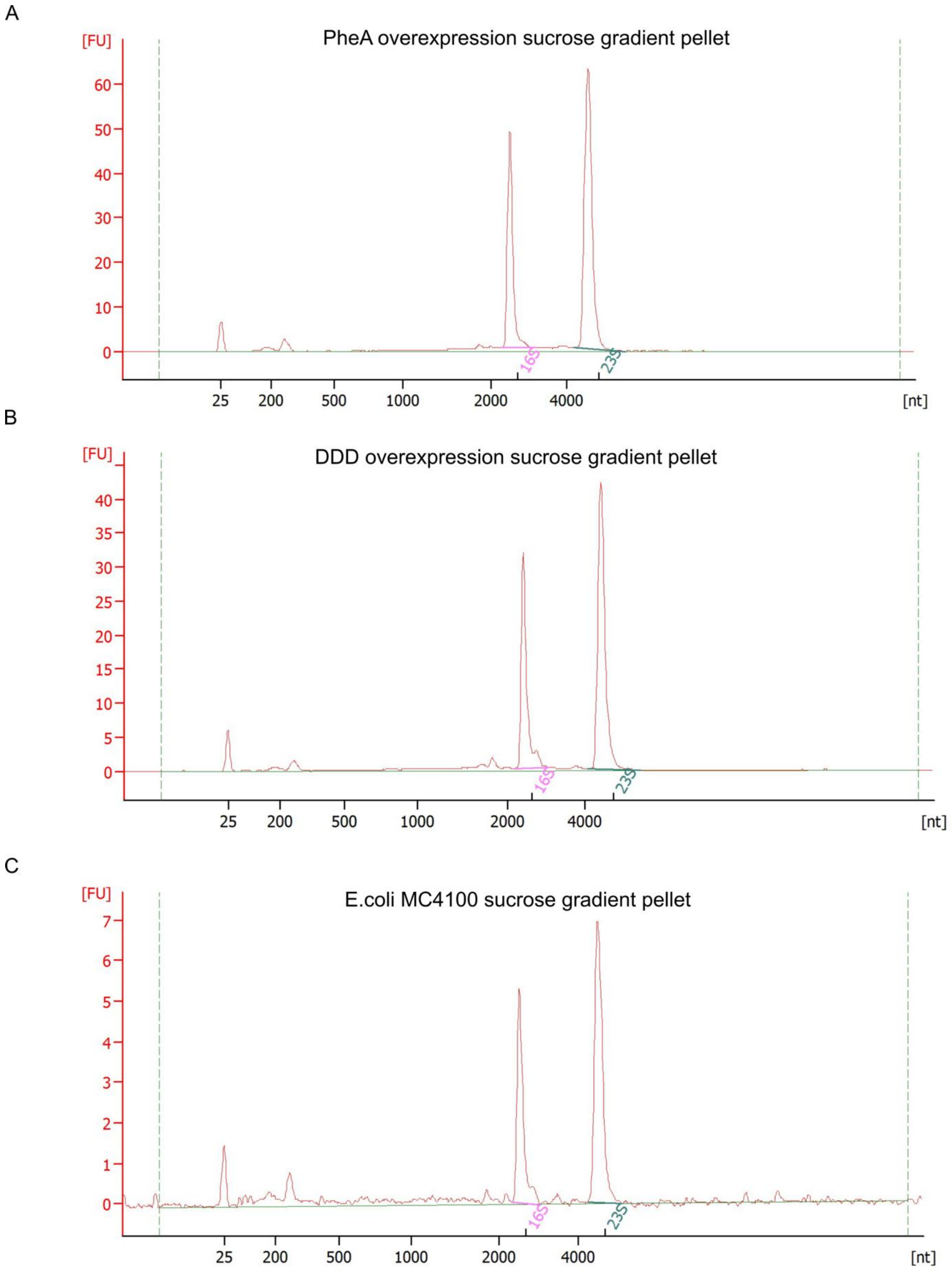


**Figure S1: RNA recovered from the pellet contains ribosomes.** Bioanalyzer Nano Chip traces of RNA recovered from the pellet of the sucrose gradient of A) wild-type, B) PheA-overexpressing, and C) DDD overexpressing cells.

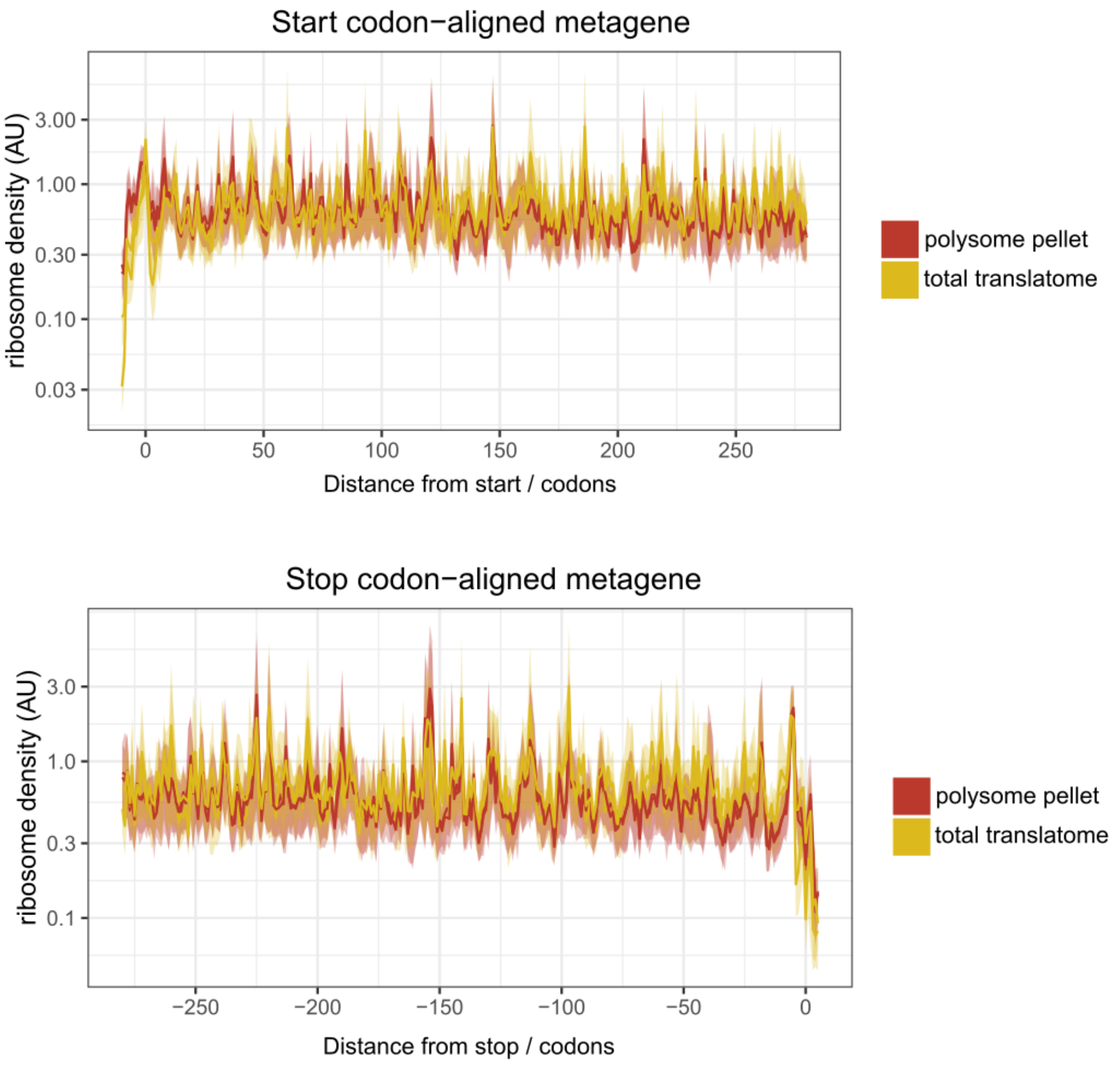


**Figure S2: Footprints of ribosomes extracted from sucrose gradient pellets resemble footprints from total translatomes.** Start (top) and stop (bottom) codon-aligned metagenes of footprints from total translatomes or pellet fractions of the sucrose gradient. Only genes longer than 300 codons are used to calculate metagenes that are at least 90 nucleotides away from another gene (n = 621).

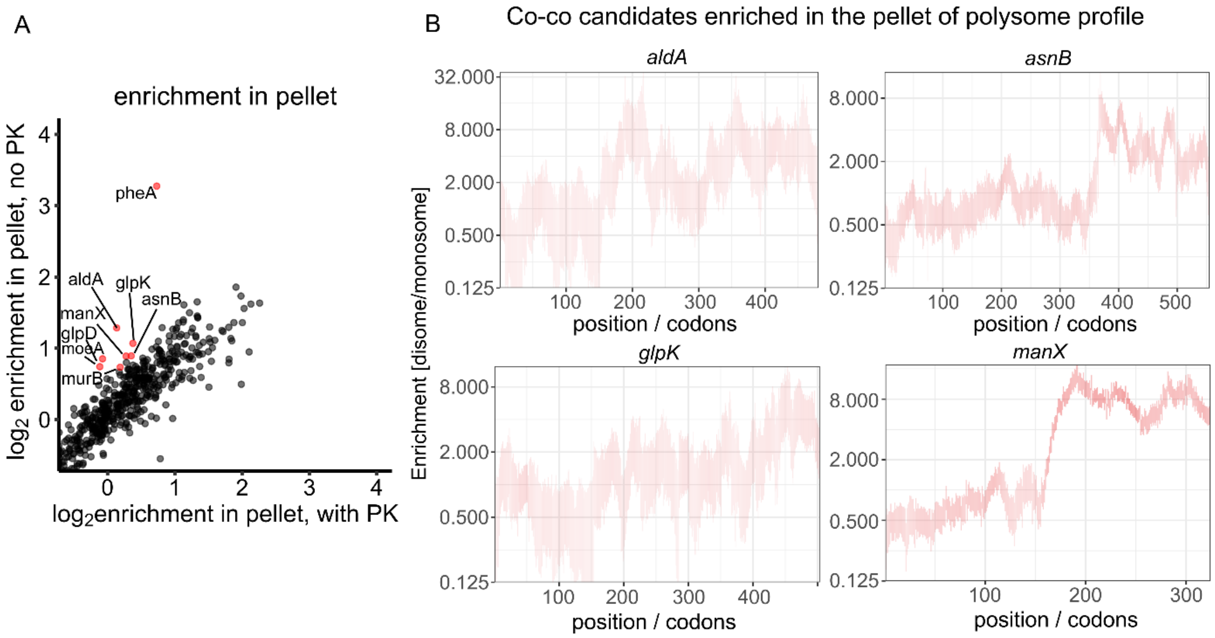


**Figure S3: Other co-co candidates may also form polysomal networks.** A) Zoom-in on the enrichment plot depicted in Figure 1G reveals several potential *trans*-assembly candidates. Each dot represents an enrichment value of ribosome protected footprints compared to total translatome in the sucrose gradient pellet without proteinase K treatment (y-axis) or after mild digestion with proteinase K (x-axis). Red color indicates at least 1.45-fold depletion in the pellet after treatment with proteinase K and at least 1.5-fold enrichment in the pellet compared to total translatome. Only genes with a mean read density of 50 reads per million (rpm) in the total translatome (out of 8 measured experiments) were plotted. B) DiSP profiles (disome/monosome) of potential co-co candidates which were also detected to be enriched in the pellet of the sucrose gradient.

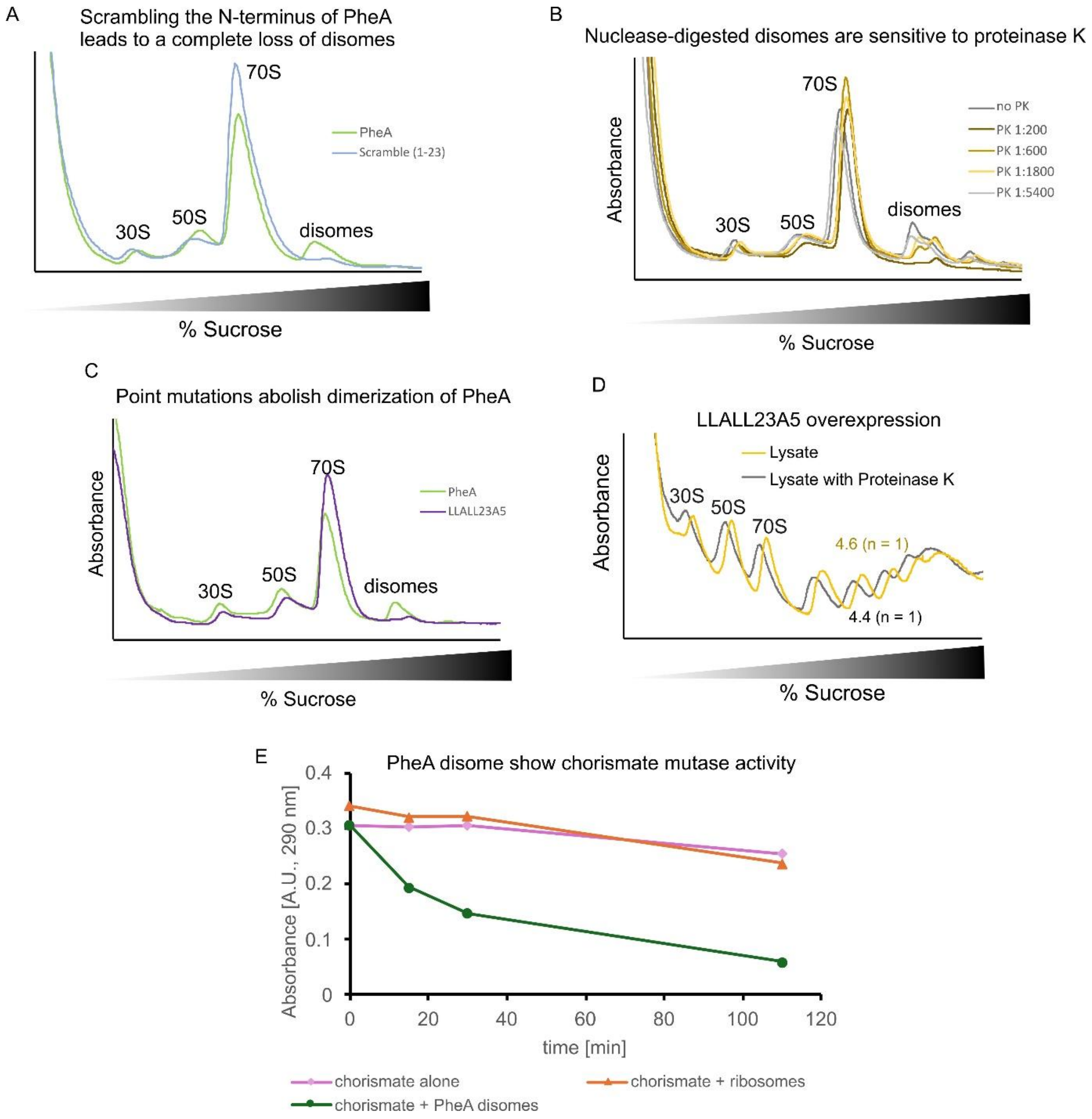


**Figure S4: Co-co assembly of PheA under overexpression conditions is native-like**. A) Polysome profiles of *E. coli* lysates overexpressing PheA wild-type (green) or PheA scramble mutant (blue) where the first 23 amino acids are randomly scrambled. B) MNase-resistant disomes are sensitive to mild proteinase K digestion. Ratios indicate the amount of added proteinase K (1 µg) per x µg of detected RNA. C) Point mutations in the dimerization domain lead to a loss of MNase-resistant disomes. D) Point mutations in the dimerization domain prevent loss of intact polysomes on sucrose gradient. E) Nuclease resistant disomes have chorismate mutase activity as detected by the decrease of absorbance at 290 nm.

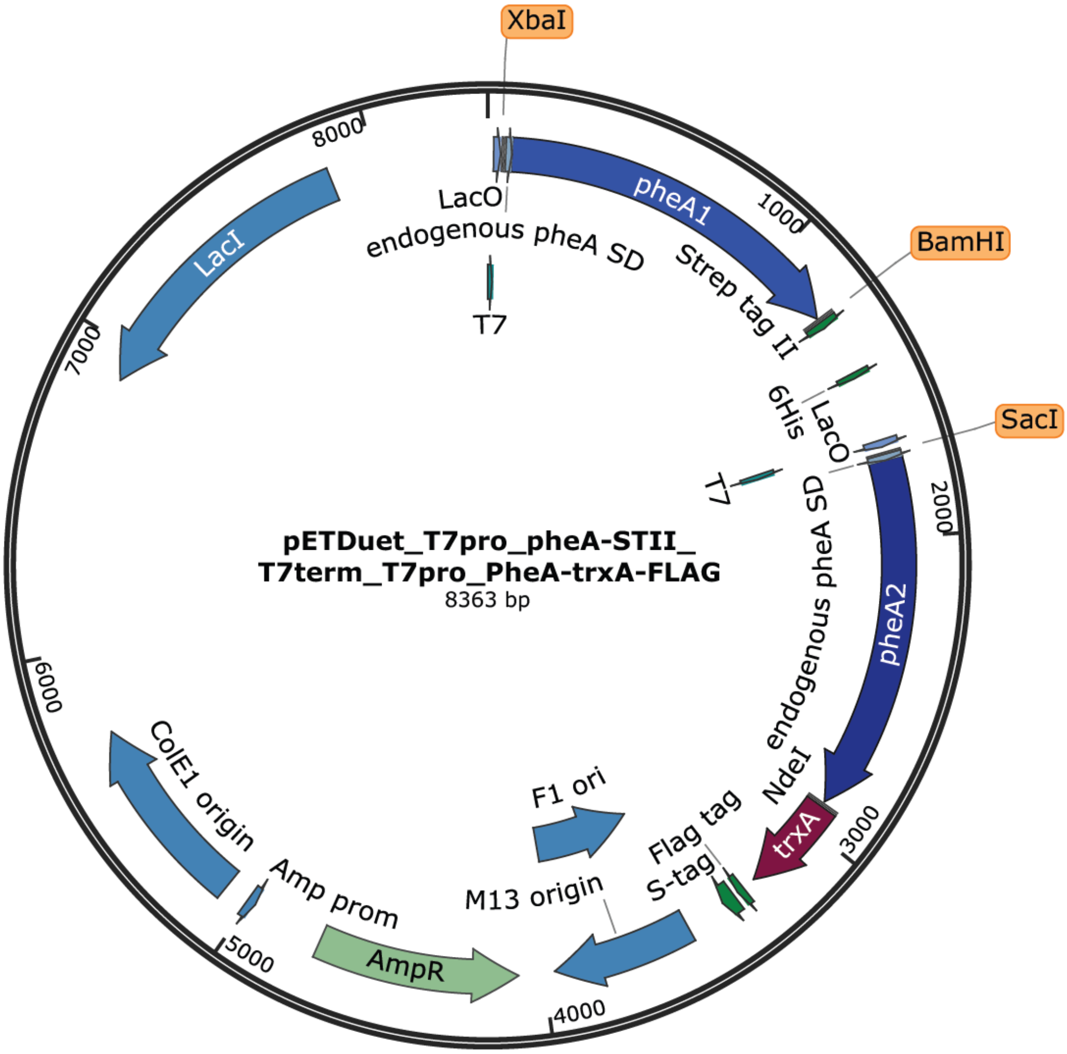


**Figure S5**: **Annotated plasmid map for 2-ORF assembly experiment.** Variants of PheA (*pheA1-strepTagII* and *pheA2-trxA-FLAG-tag*) are cloned int tandem in pETDuet vector, each under its own T7 promotor and terminator.

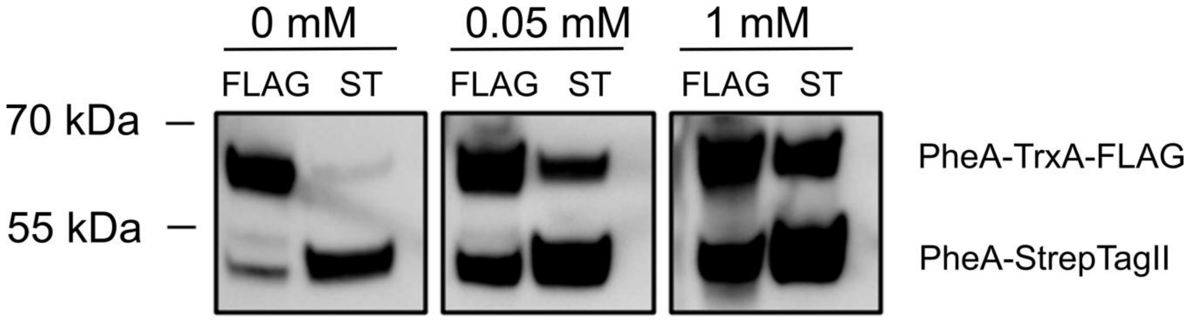


**Figure S6: PheA-TrxA-FLAG reproducibly copurifies with PheA-FLAG, results of the second replicate.** PheA-StrepTagII and PheA-TrxA-FLAG tag were cloned into pET duet vector under independent promoters and cells were induced with 0 mM, 0.05 mM, or 1 mM IPTG. Proteins were pulled on from the lysate by either using Streptactin beads (ST) or anti-FLAG beads (Flag). In each condition, we observe the co-purification of the other protein variant. 0.5, 0.125, and 0.03 volumes were loaded for 0 mM, 0.05 mM, and 1 mM sample, respectively.

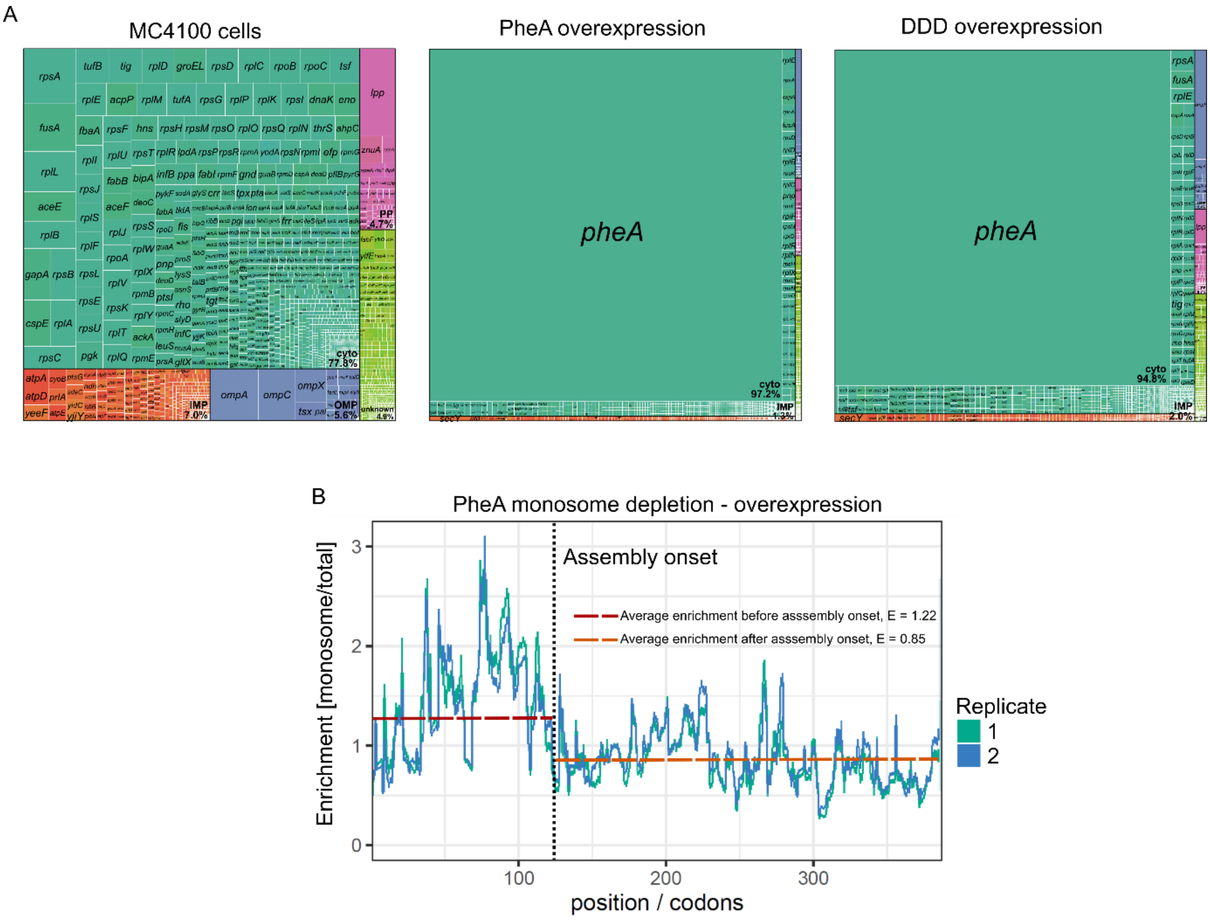


**Figure S7: Overexpression of PheA leads to the majority of ribosomes translating PheA.** A) Treemap plots of overall expression profiles of wild-type (MC4100) cells (left), PheA overexpressing cells (middle) and DDD overexpressing cells (right). B) Monosome depletion profile in the *pheA* transcript of PheA overexpressing cells. The average monosome enrichment (monosome/total) before and after assembly onset is indicated by the red and orange dashed lines, respectively.

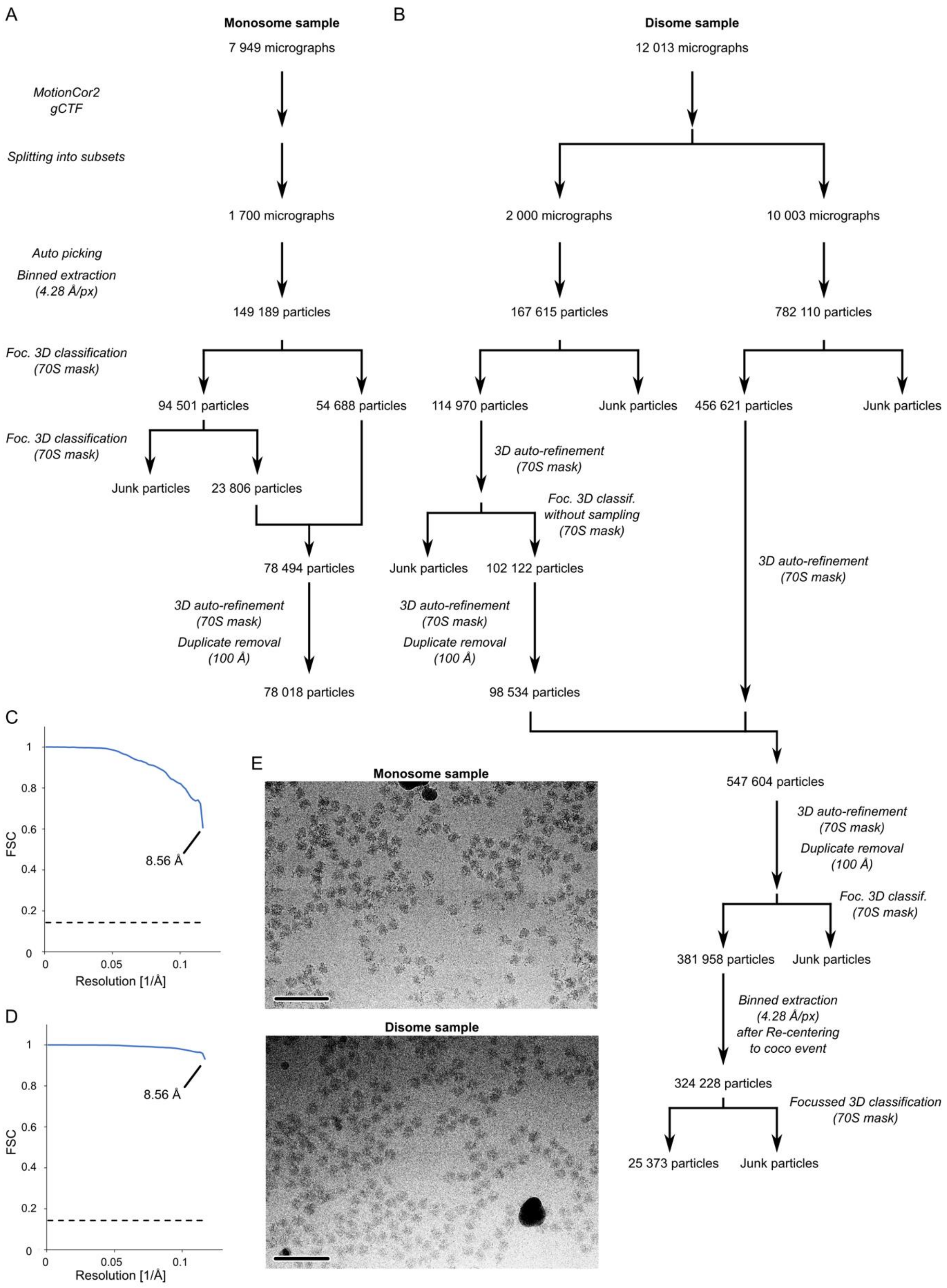


**Figure S8: Cryo-EM SPA data processing scheme for MNase-digested monosomes and disomes**. A and B) Two datasets were acquired and separately processed in Relion 3.1. Motion-corrected micrographs were split into subsets of 1700 and 2000 micrographs for monosomes (A) and disomes (B), respectively. Autopicked particles were extracted with 4x binning and iteratively classified to obtain a subset of 70S particles. After removing duplicates, TtT distances were measured. Later, the remaining disome micrographs (~ 10000) were processed in a similar manner. 70S particles of the two disome subsets were merged and classified again. The resulting 70S disome particles were further processed by the NND algorithm. C), D) The mask-corrected Fourier shell correlation (FSC) of the final reconstruction based on the merged datasets was calculated between two independently refined half-sets of the data using Relion 3.1 (FSC threshold at 0.143). E) Representative micrographs of both datasets. Scale bar: 100 nm (lower left corner).

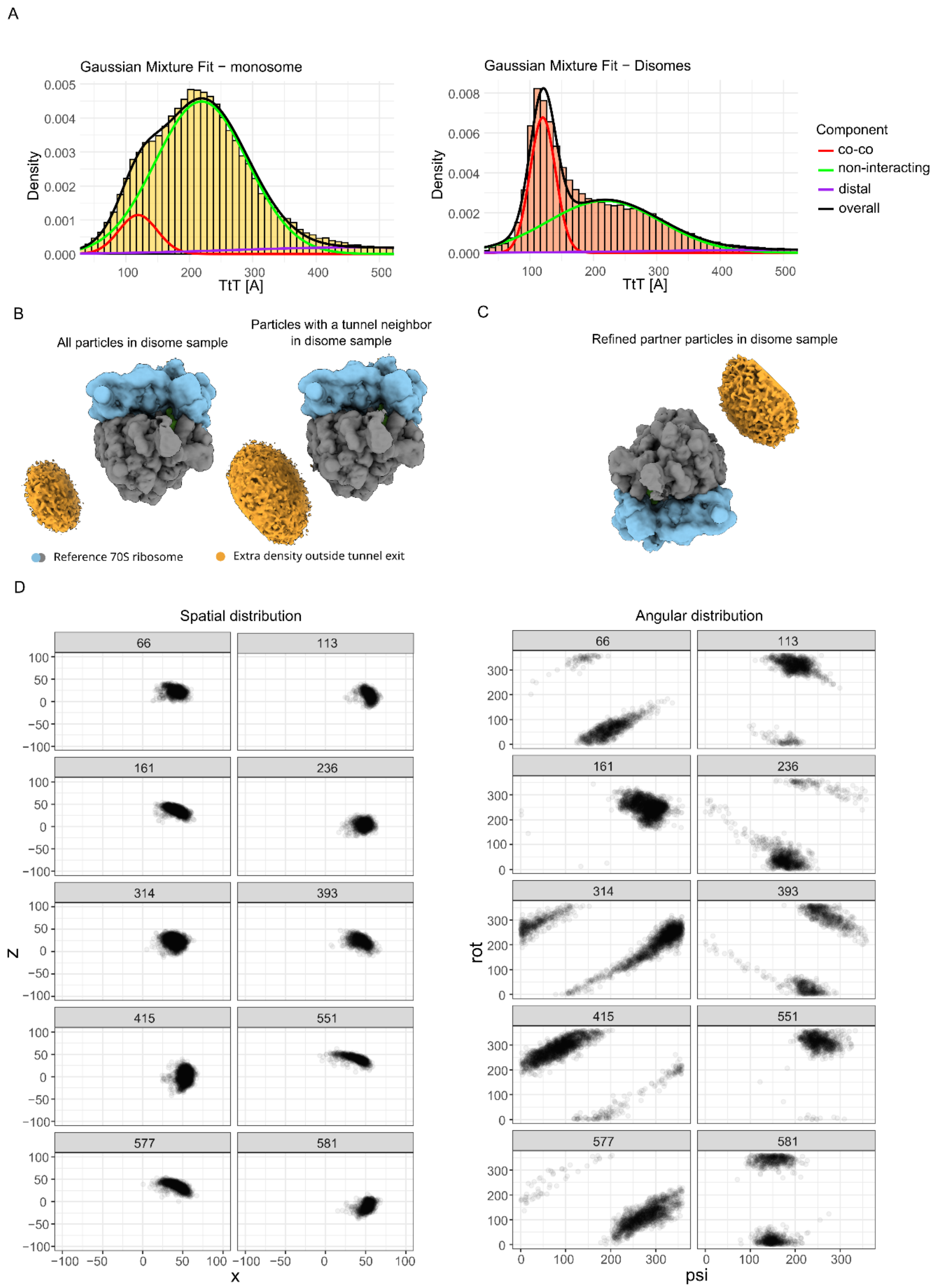


**Figure S9: Ribosomes in co-co assembly disomes are spatially unrestricted.** A) Gaussian fits of TtT distance distributions for monosome and disome samples. For each component, the mean and variance were fitted globally across both samples. B) Selecting ribosomes with a close TtT neighbour (TtT < 140 Å) leads to a more pronounced additional density at the polypeptide exit tunnel. C) Focused 3D auto-refinement on the additional density near the polypeptide exit tunnel. For this purpose, extraction coordinates of the initial particles were shifted by 250 Å in the direction of the polypeptide exit tunnel. D) Spatial and angular analysis of the 10 most abundant NND clusters accounting for 50 % of total particles of the disome dataset (15824 out of 31773). Only two dimensions are shown for each plot.

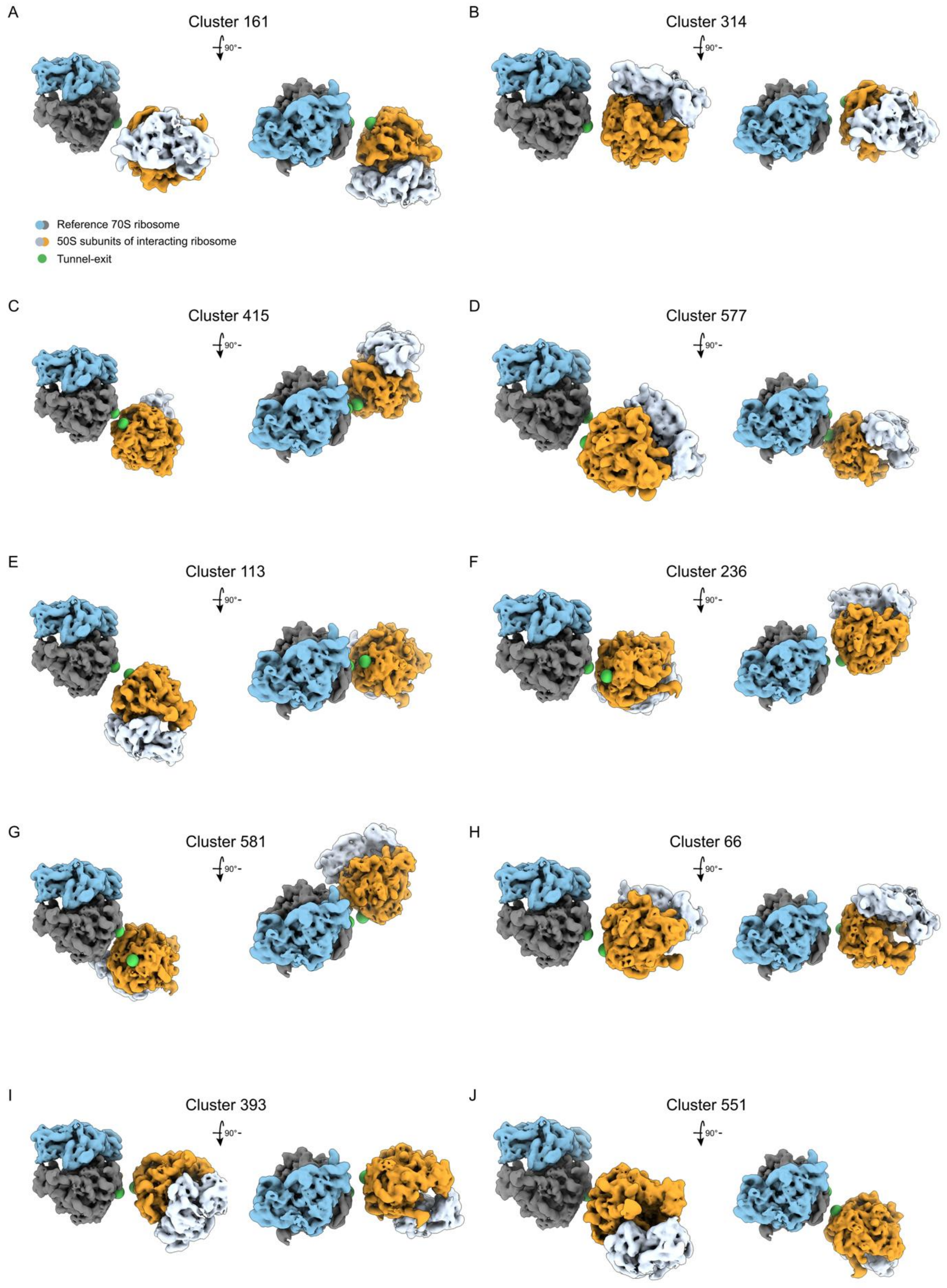


**Figure S10: Visualization of disome configurations detected by Next Neighbour Distribution-clustering.** Visualization of the clusters shown in **Figure S9D**. Reference 70S ribosomes are coloured in blue and grey, while the interacting ribosomes are coloured in orange and white. Tunnel exits are indicated by green spheres.

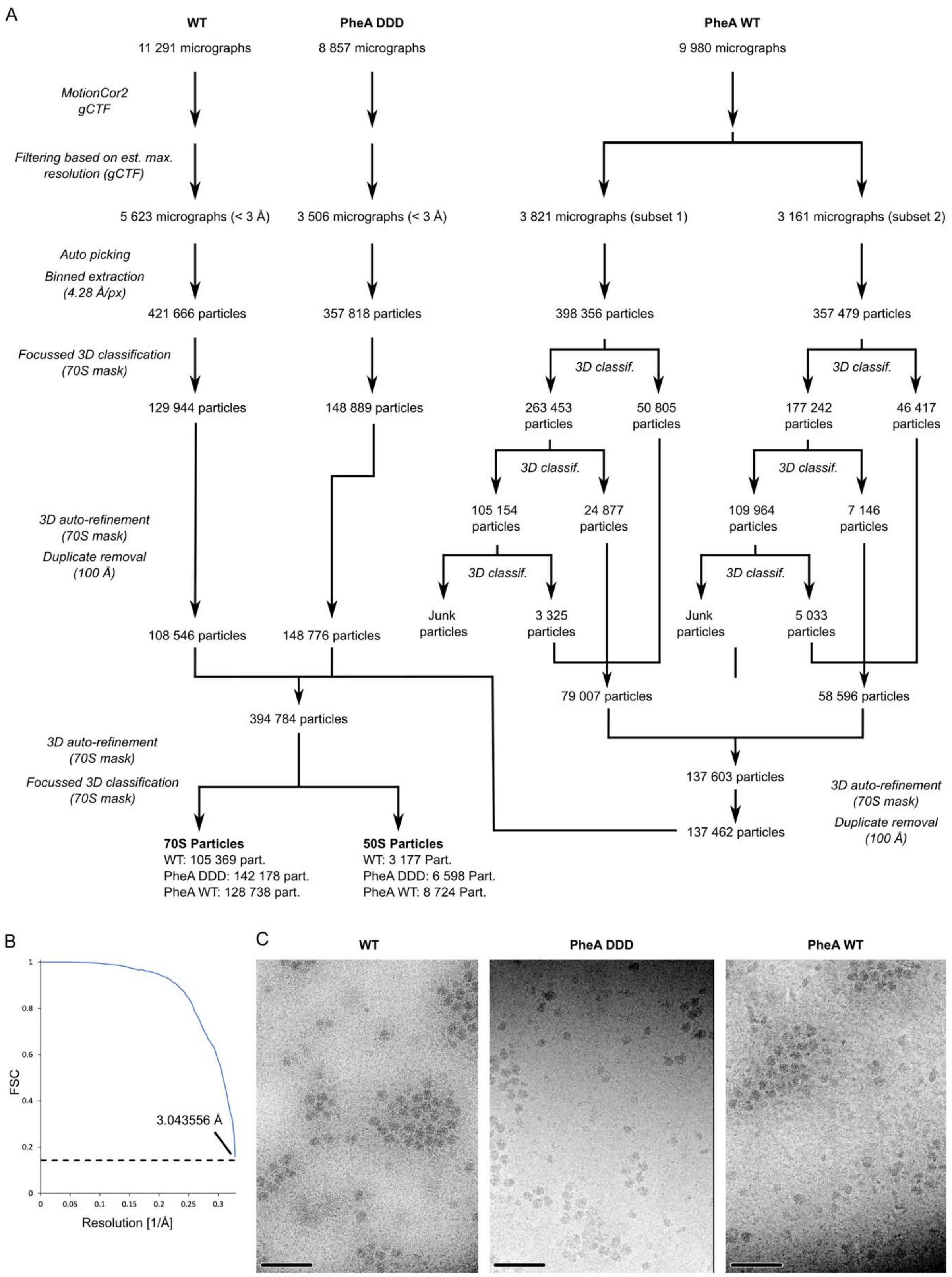


**Figure S11: Cryo-EM SPA processing scheme**. A) Three datasets were acquired and separately processed in Relion 3.1. Motion-corrected micrographs were filtered based on their maximal resolution as estimated by gCTF. Autopicked particles were extracted with 4x binning and iteratively classified to obtain a subset of true-positive particles. After removing duplicates, 70S particles were merged and refined together. B) Final cryo-EM SPA reconstruction from the merged 70S particle set of at 3.1 Å resolution. C) Mask-corrected Fourier shell correlation (FSC) of the final reconstruction (merged datasets) was calculated between two independently refined half-sets of the data using Relion 3.1 (FSC threshold at 0.143). D) Representative micrographs of the respective datasets. Scale bar: 100 nm (left lower corner)

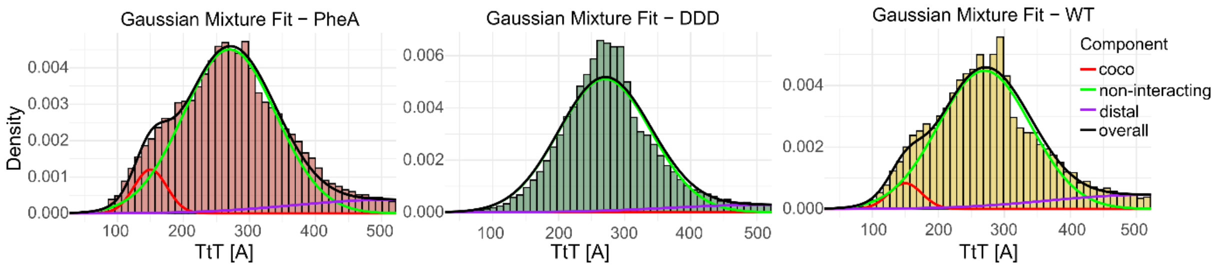


**Figure S12: Polypeptide exit tunnel proximity is preserved in intact polysomes.** Gaussian fitting to TtT distance distributions of PheA (red), DDD (green), and WT (yellow) samples.

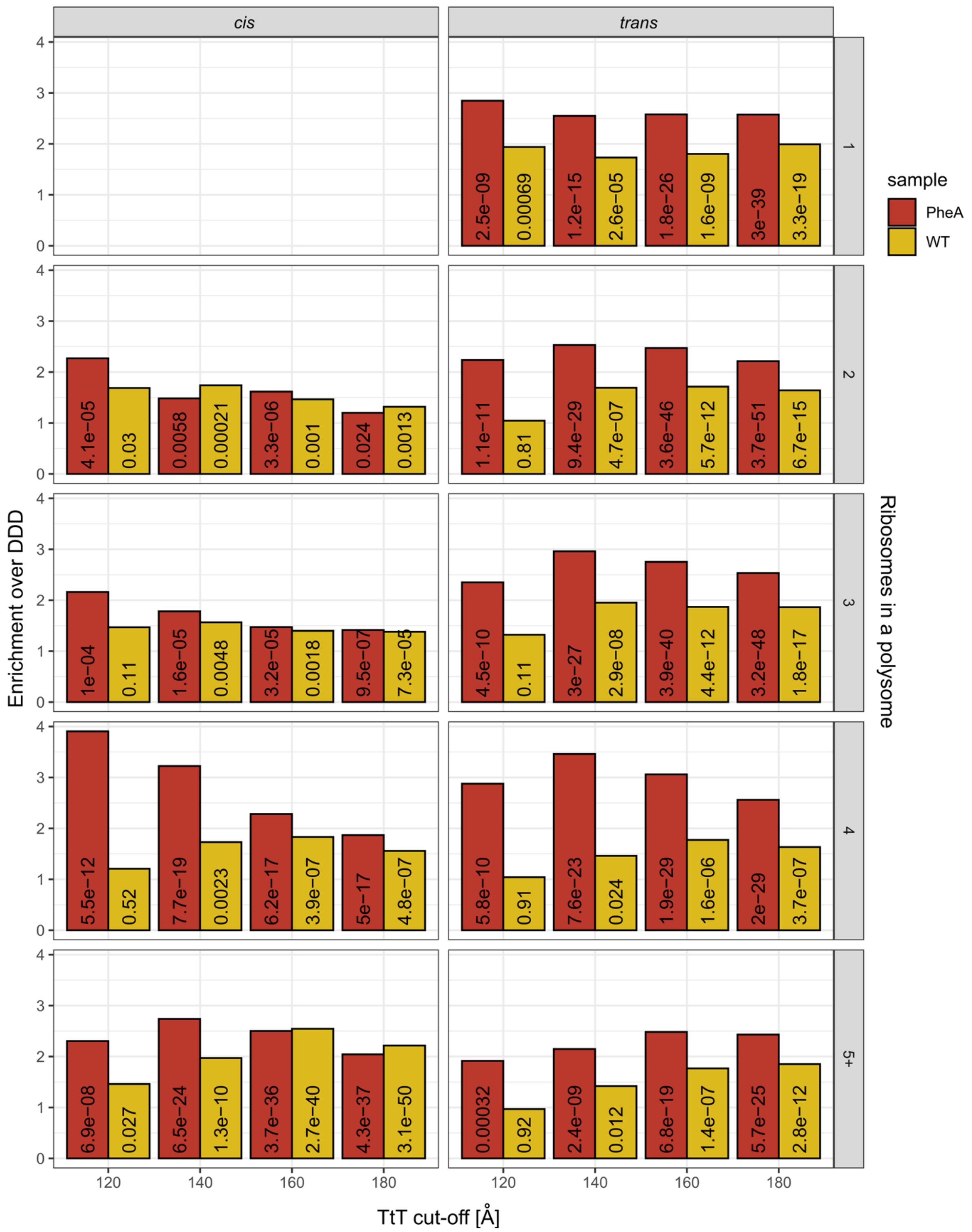


**Figure S13: Sensitivity analysis for different TtT cut-off values (cryo-EM SPA).** TtT cut-off value has only minor impact on observed enrichments in SPA datasets. The enrichments are plotted for each interaction type (left: *cis;* right: *trans*) and polysome size separately. Numbers indicate unadjusted p-value of Fisher's exact test.

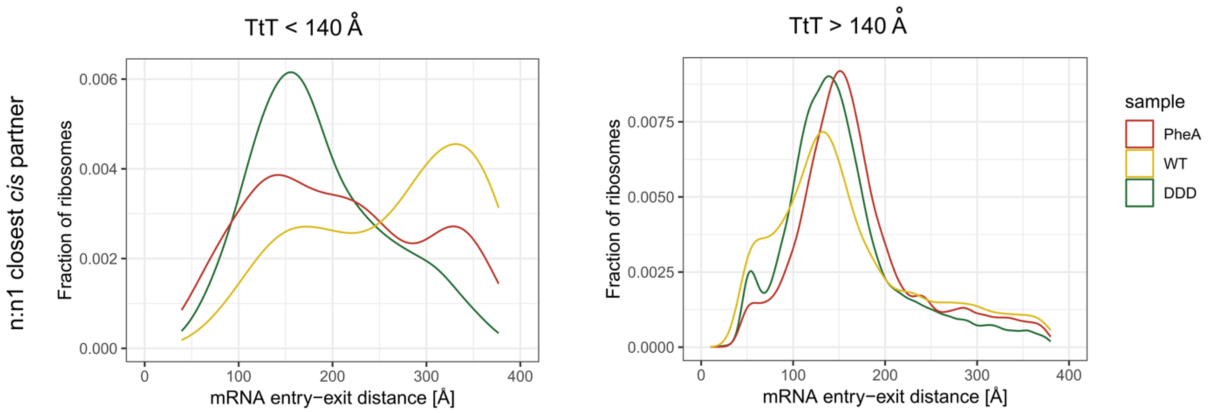


**Figure S14: Direct polysomal neighbours need longer spacing on mRNA to have short TtT distances between one another**. Above: mRNA entry-to-exit distance between direct polysomal neighbours. Left: Ribosomes within co-co distance. Right: Above the co-co distance.

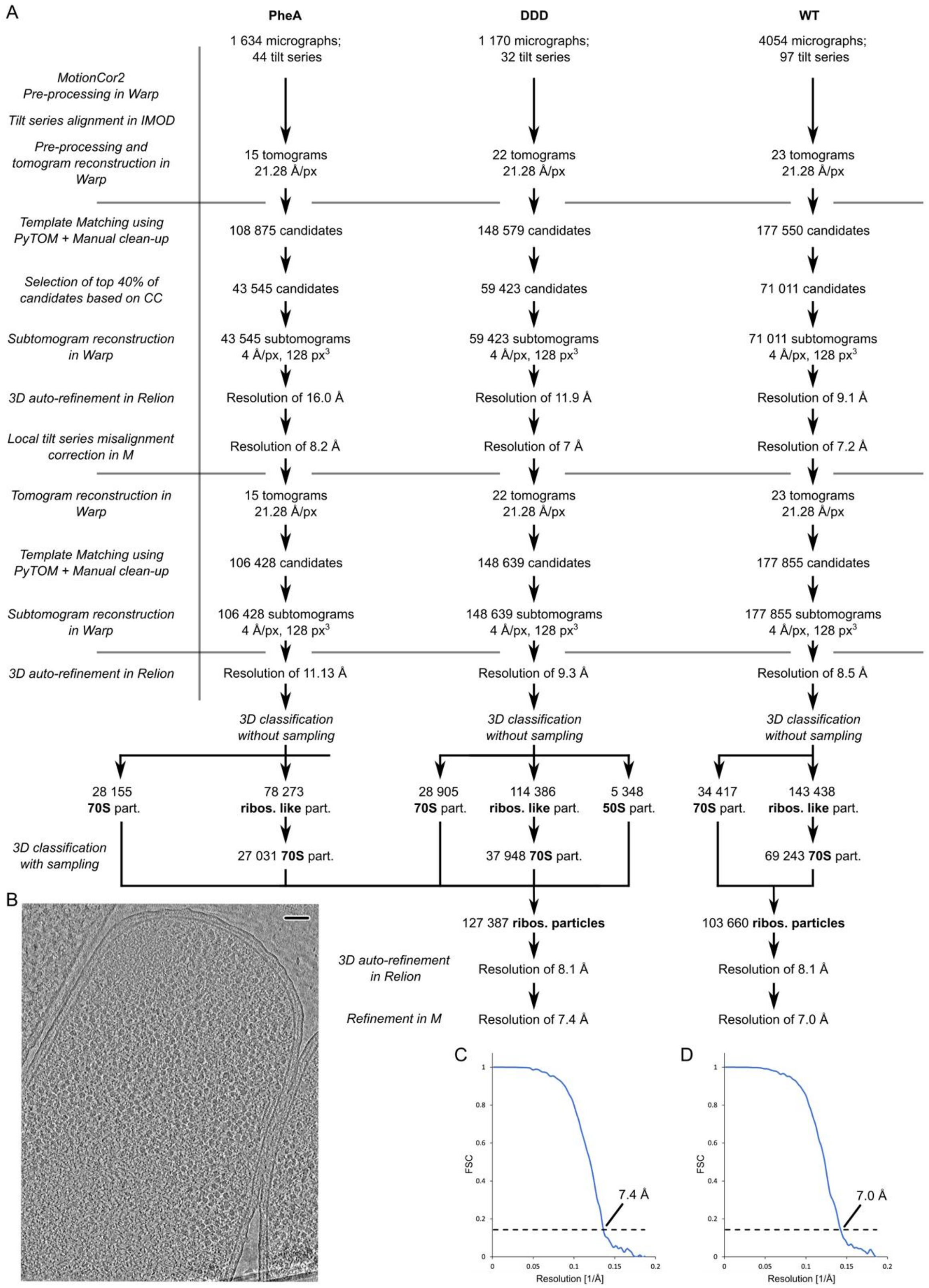


**Figure S15: Cryo-ET data processing scheme.** A) Three datasets were acquired and processed separately using the same workflow. Tilt series were pre-processed using MotionCor2 and Warp prior to tilt series alignment and tomogram reconstruction. The same reference and mask were used to localise 70S in both datasets. After optimization of tilt series alignment in M, 70S ribosomes were comprehensively localised and subjected to 3D auto-refinement. B) Z-slice through the same tomogram of PheA-overexpressing cells shown in **Figure 4A**. Scale bar: 100 nm (Upper right corner). C and D) Mask-corrected gold-standard Fourier shell correlation (FSC) was calculated between two independently refined half-sets of the data using M (FSC threshold at 0.143). C) Merged consensus reconstruction of the PheA and DDD datasets. D) Merged consensus reconstruction of the WT dataset.

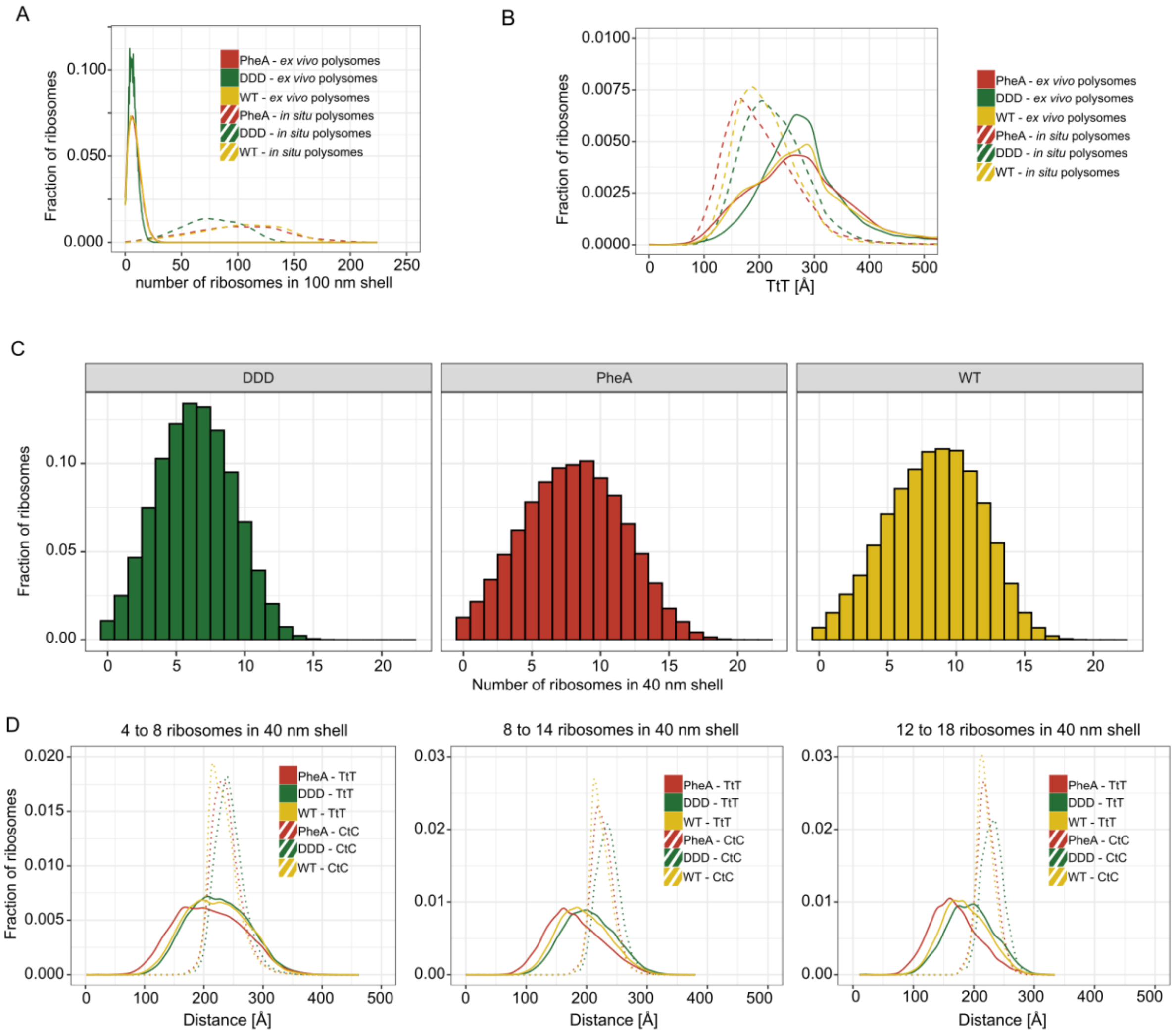


**Figure S16: Polypeptide exit tunnel proximity is preserved *in situ* regardless of local ribosome density.** A) Overall ribosome counts in 100 nm shell in *ex vivo* polysomes analyzed by cryo-EM SPA (**Figure 3**) and *in situ* polysomes analyzed by tomography (**Figure 4**). B) TtT distance distributions of *ex vivo* polysomes and *in situ* polysomes. C) Local ribosome density characterized by the number of ribosomes found in a 40 nm shell. D) Varying local ribosome density does not impact overall TtT distance distributions. By restricting local density, we observed that the distribution of distances between centers of ribosomes (center-to-center, CtC) were similar between the samples, indicating reorientation of ribosomes during co-co assembly.

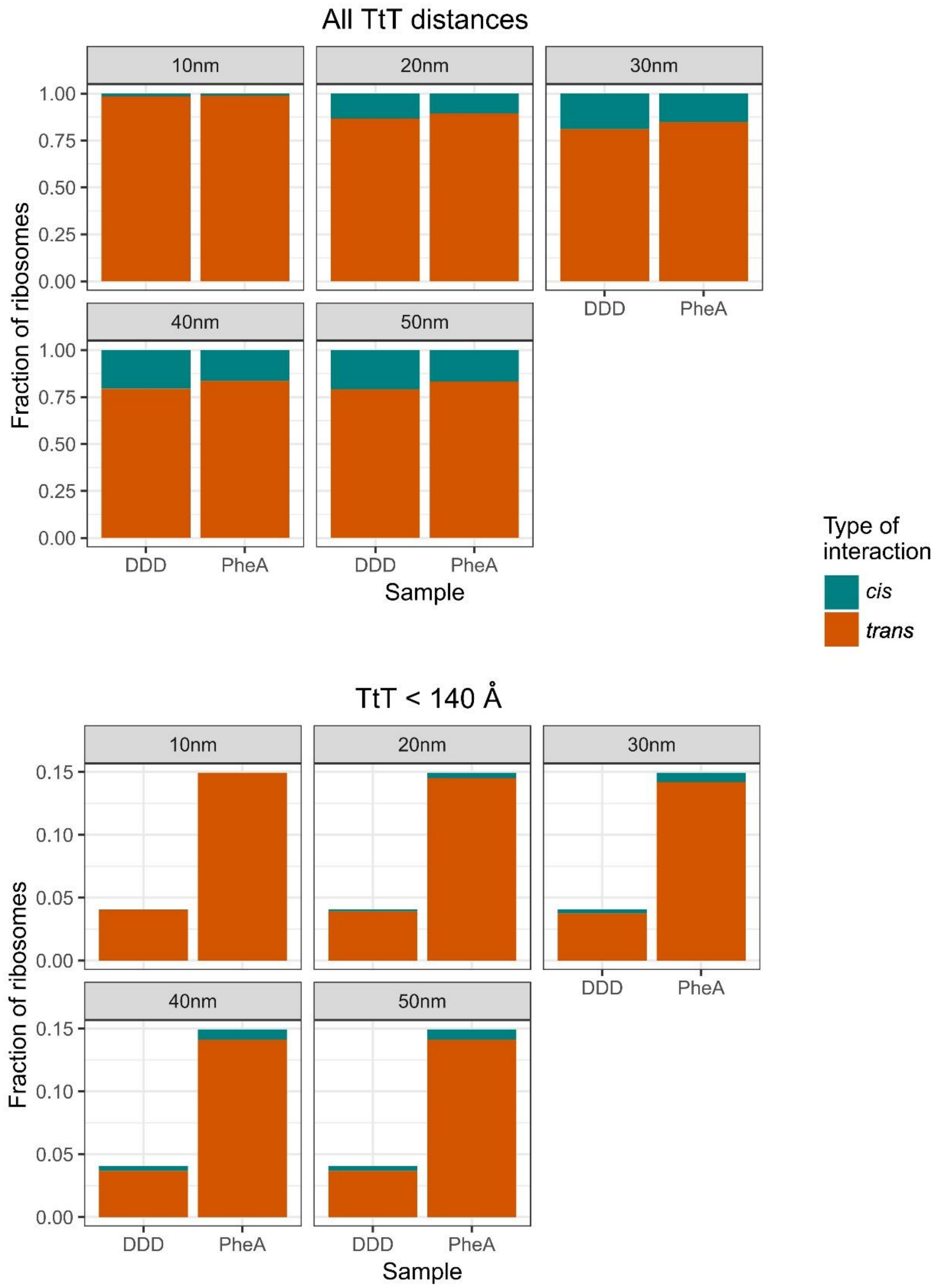


**Figure S17: mRNA entry-exit search range beyond 20 nm value has little impact on cis/trans distribution.** Assignment of closest TtT neighbours on either same mRNA (*cis*) or different mRNA (*trans*) across 10, 20, 30, 40, and 50 nm polysomal neighbour search ranges and all distances (top) or co-co distances (bottom)

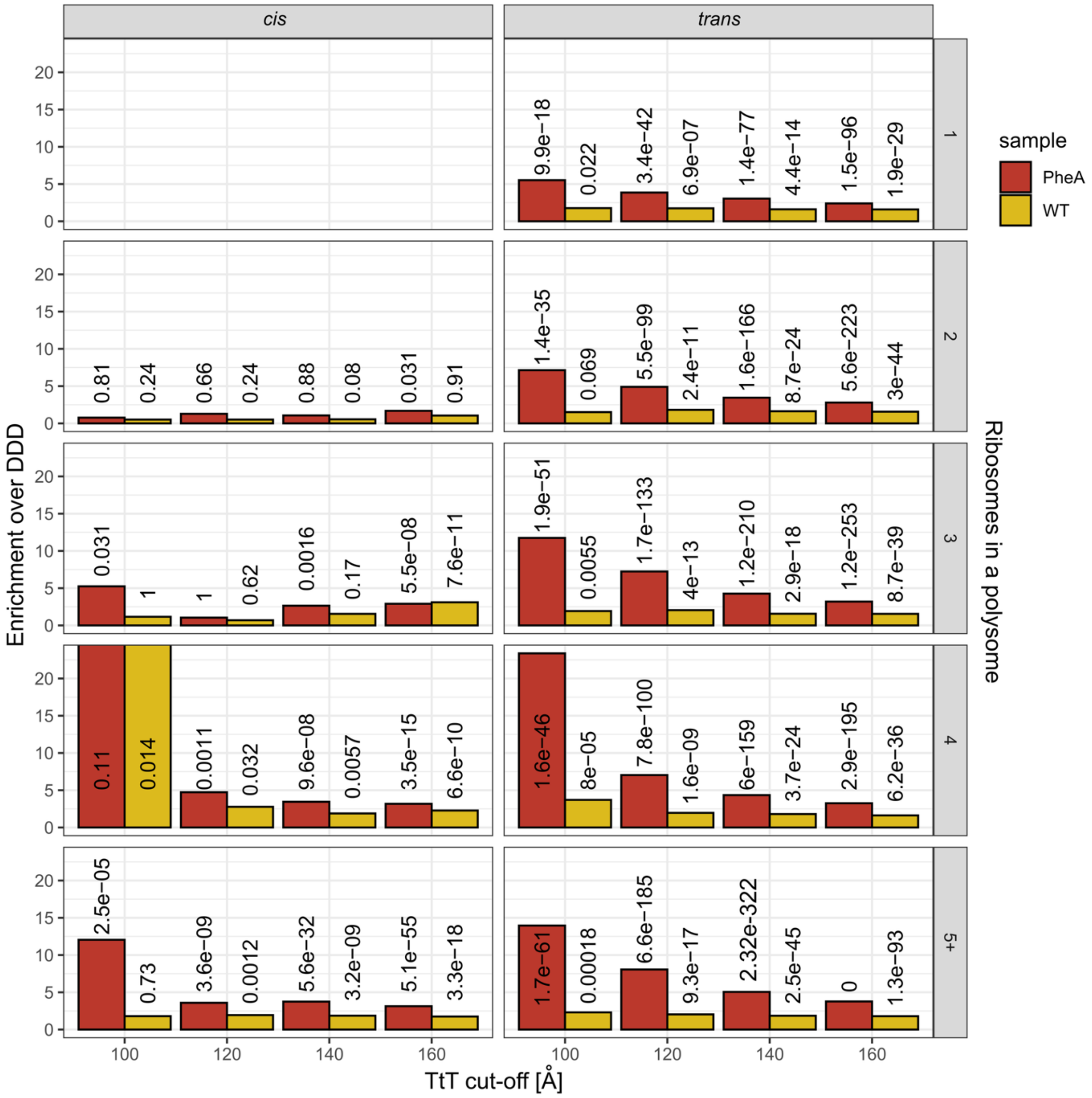


**Figure S18: Sensitivity analysis for different TtT cut-off values (cryo-ET).** Selection of different TtT distance cut-off has little impact on overall results of ribosome enrichment in the tomography data. The enrichments are plotted for each interaction type and polysome size separately. Number indicate unadjusted p-value of Fisher's exact test.

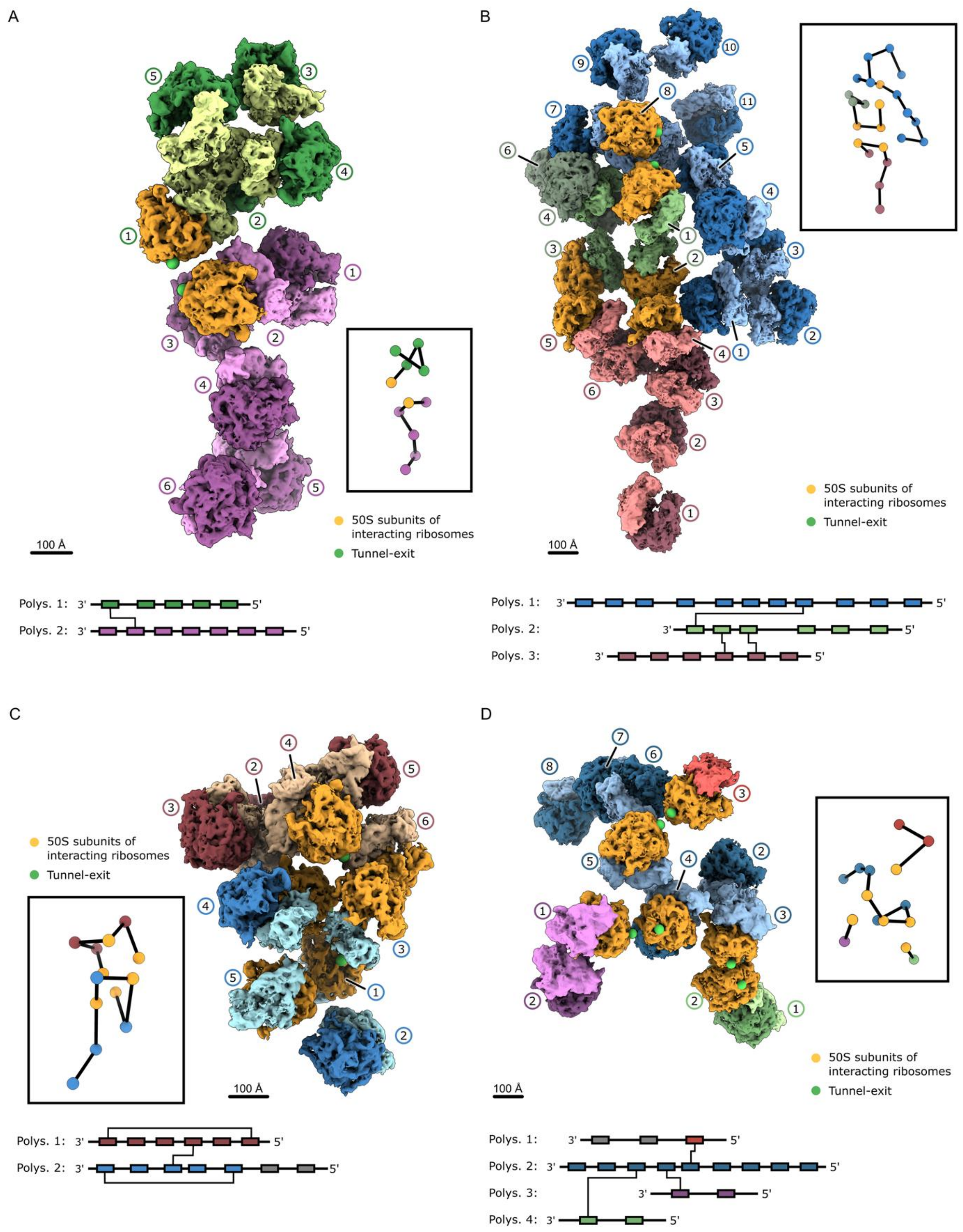


**Figure S19: 3D visualization of TtT proximities in *trans***: A-D) Individual polysomes are coloured differently. The 50S subunits of *cis*- or *trans*- interacting ribosomes are highlighted in orange; the tunnel exits by green spheres. TtT interactions between polysomes are schematically illustrated below the 3D visualization. The center-to-center traces of polysomes are shown in boxes.

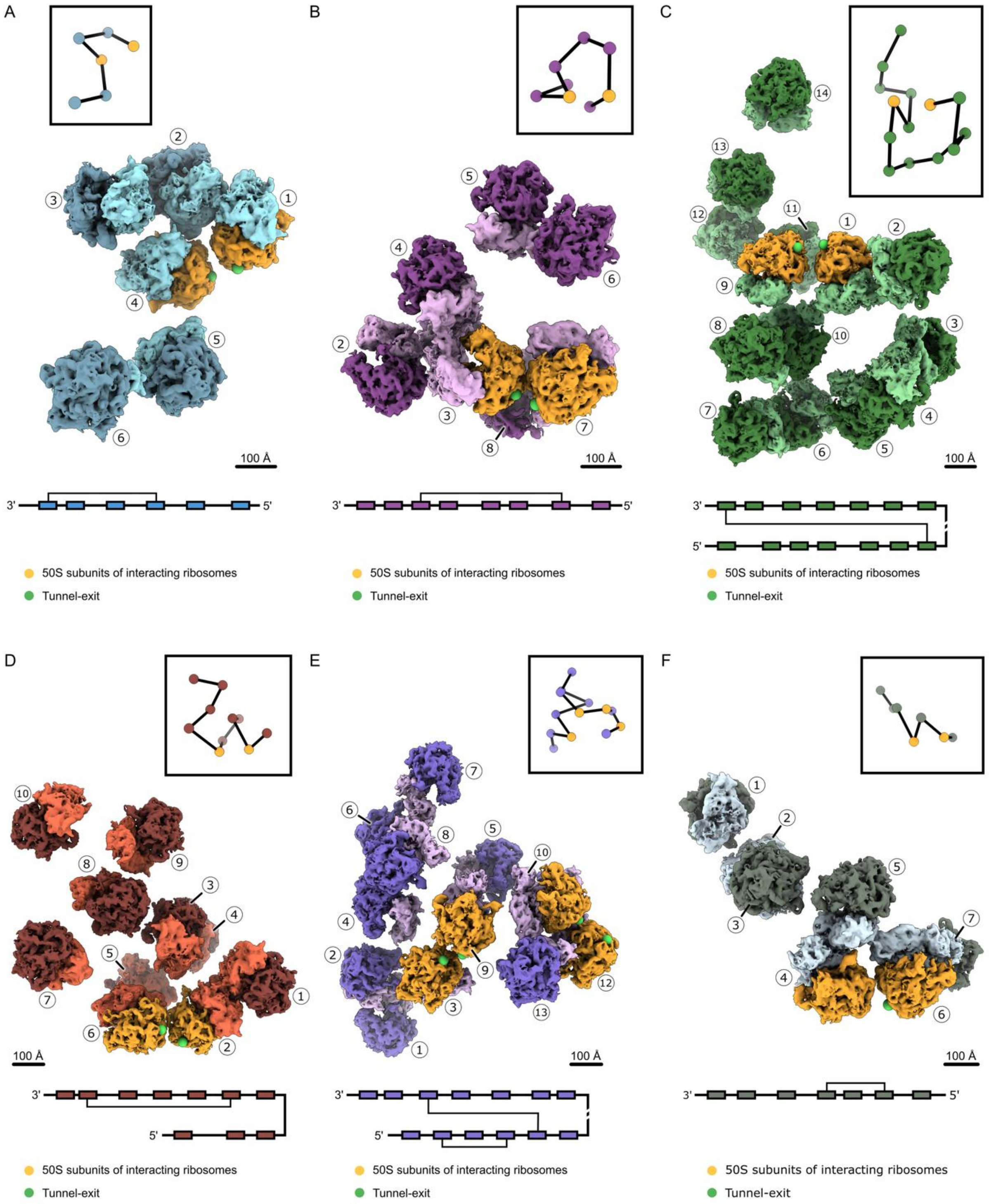


**Figure S20: 3D visualization of TtT proximities in *cis***: A-F) The 50S subunits of *cis*-interacting ribosomes are highlighted in orange; the tunnel exits by green spheres. TtT interactions within the polysomes are schematically illustrated below the 3D visualization. The center-to-center traces of polysomes are shown in boxes.

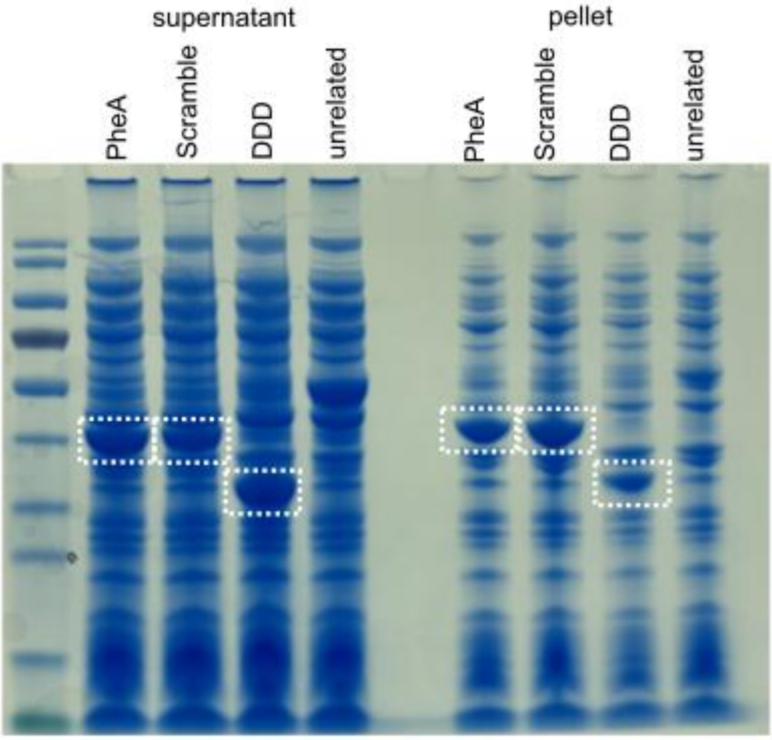


**Figure S21: The dimerization domain is the major cause of PheA aggregation.** Coomassie blue stained protein SDS-page of *E. coli* overexpressing different proteins. The lysate was divided into supernatant and pellet prior to loading. The band of the overexpressed protein is highlighted by a white box.

**Supplementary Table 1:** Cryo-EM SPA data collection and refinement statistics

| **Data collection and processing** | **WT** | **DDD** | **PheA** | **Monosomes** | **Disomes** |
|---|---|---|---|---|---|
| Magnification | 81 k | 81 k | 81 k | 81 k | 81 k |
| Voltage (kV) | 300 kV | 300 kV | 300 kV | 300 kV | 300 kV |
| Electron exposure (e–/Å$^2$) | 45.26 | 45.0 | 43.6 | 45.5 | 45.5 |
| Defocus range (µm) | -0.75 to -1.75 | -0.75 to -1.75 | -0.75 to -1.75 | -0.75 to -1.75 | -0.75 to -1.75 |
| Pixel size (Å) | 1.07 | 1.07 | 1.07 | 1.07 | 1.07 |
| Symmetry imposed | C1 | C1 | C1 | C1 | C1 |
| Final particle images (no.) | 105 369 | 142 178 | 128 738 | 78 018 | 25 373 |
| Map resolution (Å) | | 3.04 | | 8.56 | 8.56 |
| FSC threshold | | 0.143 | | 0.143 | 0.143 |
| Map resolution range (Å) | | 3.04 - 6 Å | | 8.56 – 11 Å | 8.56 – 11 Å |

**Supplementary Table 2:** Cryo-ET data collection and refinement statistics

| **Data collection and processing** | **DDD** | **PheA** | **WT** |
|---|---|---|---|
| Magnification | 33 k | 33 k | 33 k |
| Voltage (kV) | 300 kV | 300 kV | 300 kV |
| Electron exposure per frame (e–/Å$^2$) | 3.9 | 3.78 | 3.8 |
| Defocus range (µm) | -3.0 to -6.0 | -3.0 to -6.0 | -3.0 to -6.0 |
| Pixel size (Å) | 2.54 | 2.54 | 2.54 |
| Symmetry imposed | C1 | C1 | C1 |
| Final particle images (no.) | 66 853 | 55 186 | 103 660 |
| Map resolution (Å) | 7.4 | | 7.0 |
| FSC threshold | 0.143 | | 0.143 |
| Map resolution range (Å) | 5.08 - 10 | | 5.08 - 10 |
